# Methane formation driven by light and heat prior to the origin of life

**DOI:** 10.1101/2023.04.11.535677

**Authors:** Leonard Ernst, Uladzimir Barayeu, Jonas Hädeler, Tobias P. Dick, Judith M. Klatt, Frank Keppler, Johannes G. Rebelein

## Abstract

Methane is a potent greenhouse gas, which likely enabled the evolution of life by keeping the early Earth warm. Here, we demonstrate new routes towards abiotic methane formation under early-earth conditions from methylated sulfur and nitrogen compounds with prebiotic origin. These compounds are demethylated in Fenton reactions governed by ferrous iron and reactive oxygen species, produced by light and heat in aqueous environments. The reactions generate methyl radicals and ultimately release methane and ethane. Organic iron chelators enhance reaction rates and recycle ferric to ferrous complexes via ligand-to-metal charge transfer, establishing a light-driven iron redox cycle. This abiotic reaction facilitates methane and ethane formation across Earth’s humid realm, thereby shaping the chemical evolution of the atmosphere prior to the origin of life and beyond.

**One-Sentence Summary:** Under suboxic and anoxic conditions, iron and reactive oxygen species drive the global formation of methane in aqueous environments.

## Main Text

Methane (CH4) is a potent greenhouse gas which has in the past and is still today contributing to climate change(*1*). Atmospherically accumulated CH_4_ might also explain the “faint young sun paradox”, which describes the apparent contradiction of a fainter sun (70% – 83% of the current solar energy output) but a climate that was at least as warm as today during early Earth (4.5–2.5 Ga ago)(*2*–*4*). Although these CH_4_ levels would be essential to keep the Earth a liquid hydrosphere to allow the evolution of life during the Archean (4.0–2.5 Ga), the source of CH_4_ prior to the origin of life is still under debate(*5*). While CH_4_ was released by submarine volcanism, most CH_4_ is suggested to be formed as side product of serpentinization(*5*). After the evolution of microbial methanogenesis latest by 3.5 Ga(*6*), methanogenesis could have been responsible for a CH_4_ flux comparable to today(*7*). Thus, methanogenesis is expected to be the main source of CH_4_ during the Archean, supported by light carbon isotope values in sedimentary deposits(*8*). However, isotope signals can only manifest upon reoxidation and CH_4_ itself does not leave much of a signature in the geological record. Thus, the actual CH_4_ concentrations and the potential abiotic sources during early Earth remain elusive. Based on mass-independent fractionation of sulfur, at least 20 ppmv CH_4_ was present around 2.4 Ga ago(*9*). A more recent study analyzing the fractionation of xenon isotopes suggests CH_4_ levels of >5000 ppmv around 3.5 Ga ago(*10*). Catling *et al.* expect even higher CH_4_ levels at the beginning of the Archean (4 Ga)(*3*) before methanogenesis evolved. Yet, the processes responsible for these high CH_4_ levels and their relative contributions remain controversial.

Recently, we discovered a non-enzymatic CH_4_ formation mechanism expected to occur in all living organisms(*11*). The mechanism has been demonstrated to be active in over 30 very diverse organisms(*11*) and suggested to explain previously observed CH_4_ formation by cyanobacteria(*12*), freshwater and marine algae(*13, 14*), saprotrophic fungi(*15*) and plants(*16*). The CH_4_ formation is driven by a cascade of radical reactions, governed by the interplay of reactive oxygen species (ROS) and ferrous iron (Fe^2+^), methylated sulfur (S)- and nitrogen (N)-compounds are oxidatively demethylated by hydroxyl radicals (·OH) and oxo-iron(IV) complexes ([Fe^IV^=O]^2+^) to yield methyl radicals (·CH_3_) (*11*). We wondered, if this abiotic mechanism could also occur outside living cells and might have contributed to CH_4_ levels before life emerged. All needed components (i) methylated S- and N-compounds, (ii) Fe^2+^ and (iii) ROS are found under early-earth conditions. (i) In a prebiotic world, methylated S-compounds like methanethiol, dimethyl sulfide (DMS) or dimethyl sulfoxide (DMSO) were formed abiotically under the reducing conditions of hydrothermal vents(*17*–*19*) or transported to Earth by carbonaceous meteorites during early Earth meteorite bombardment(*20, 21*). Upon the emergence of life, more methylated S-/N-compounds were produced by cells and organisms, *i.e.* methionine, dimethylsulfoniopropionate or trimethylamine(*22*). (ii) Under the anoxic conditions of the early Earth, oceans were rather ferruginous, *i.e.* rich in Fe^2+^ required for Fenton chemistry(*23, 24*), nonetheless ferric iron (Fe^3+^) also occurred in Archean seawater(*25*). Additionally, the mechanism driven by Fe^2+^ can be enhanced by Fenton-promoting Fe^2+^-chelators, e.g. ATP or citrate(*26*). Under anoxic conditions, Fe(III)-carboxylate complexes are photochemically reduced via ligand-to-metal charge transfer (LMCT)(*27*), resulting in Fe^2+^ and organic radicals(*28*) (iii) Under ambient temperatures, low ROS levels exist in water that increase with heat(*29*), or can be generated by photolysis or radiolysis (*30–33*). Under acidic conditions, *i.e.* in volcanic lakes(*34*), illumination of Fe(III)-aqua complexes ([Fe(H_2_O)_6_]^3+^) form Fe^2+^ and ROS(*35, 36*). Thus, we hypothesized that the Fenton reaction of Fe^2+^ with H_2_O_2_, generated by heat and light, could have driven the formation of CH_4_ from methylated S-/N-compounds independent of temperatures and pressures occurring at hydrothermal vents but at ambient conditions as early as the prebiotic world of the Hadean (4.5– 4.0 Ga, Fig. 1). To identify critical components of such a mechanism, we used aqueous model systems to determine the influence of heat, light, and (bio)molecules on CH_4_ formation in abiotic and biotic environments.

**Fig. 1.**
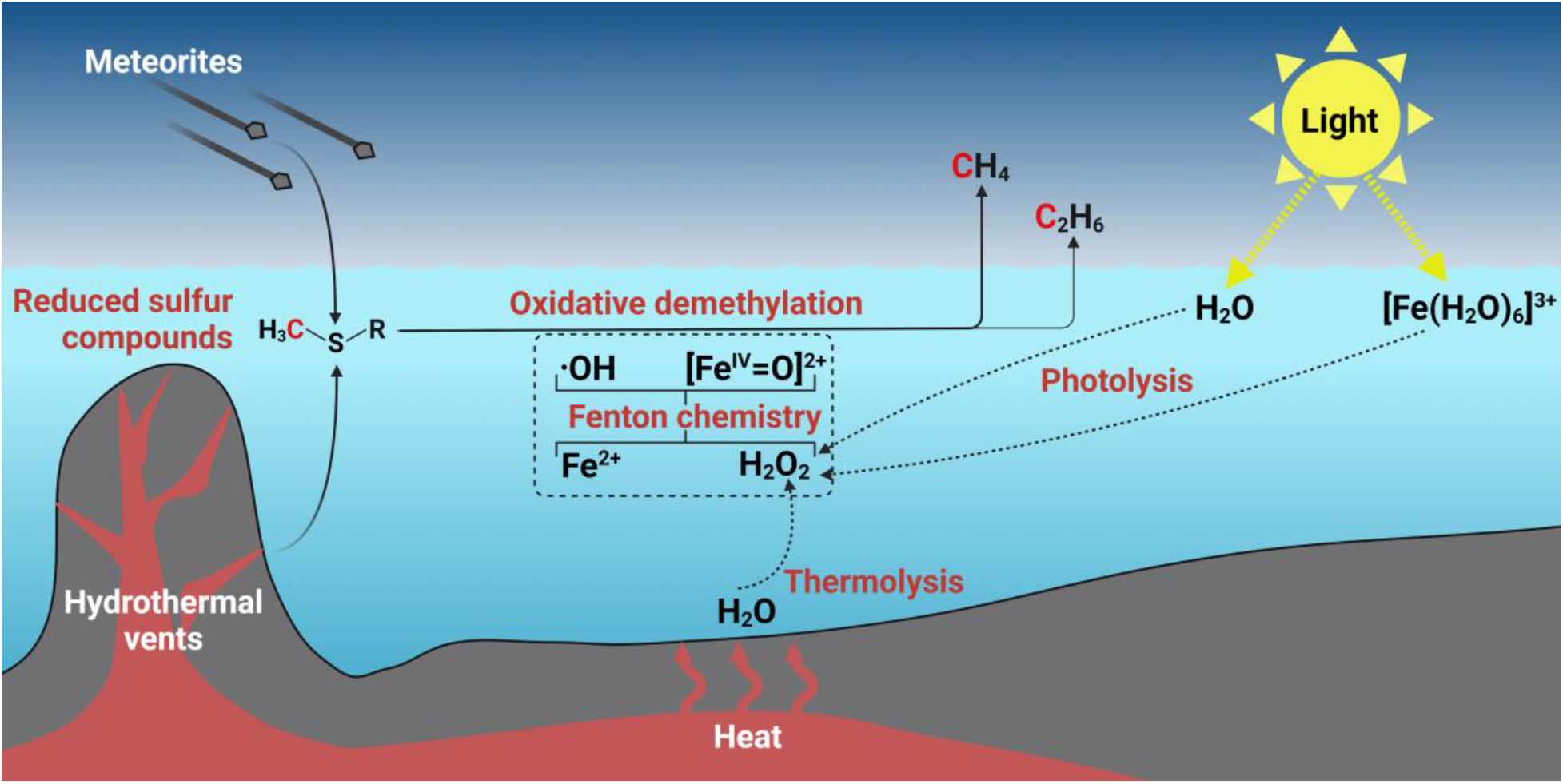
Environmental CH_4_ and C_2_H_6_ formation in a prebiotic world. Reduced, methylated S-/N-compounds are formed abiotically in hydrothermal vents or transported to Earth by carbonaceous meteorites. Under anoxic conditions, H_2_O_2_ is formed by thermolysis and photolysis of water and [Fe(H_2_O)_6_]^3+^ complexes, reacting with dissolved ferrous iron (Fe^2+^) to hydroxyl radicals (·OH) and [Fe^IV^=O]^2+^ compounds that drive the oxidative demethylation of methylated S-/N-compounds, thereby facilitating CH_4_ and C_2_H_6_ formation.

### Methane is formed under abiotic conditions

To investigate CH_4_ formation under abiotic conditions (Fig. 1), we designed a chemical model system consisting of a nitrogen atmosphere, a potassium phosphate-buffered solution (pH 7, expected during the Archean at 4.0 *Ga(37)*) supplemented with Fe^2+^ and the abiotically formed DMSO which serves as methyl donor for ROS-driven CH_4_ formation. In this model system, CH_4_ was consistently formed from DMSO in the dark (Fig. 2A). CH_4_ formation rates increased with rising temperatures from 30 °C to 97 °C, consistent with the previously reported temperature-dependency of ROS levels in water(*29*). While only marginal CH_4_ formation rates derived from DMSO were observed at 30 °C (~0.02 μM h^-1^), rates increased 41-fold to ~0.82 μM h^-1^ at 97 °C. In addition, low ethane (C_2_H_6_) amounts were formed (Fig. S1), most likely resulting from the recombination of two methyl radicals(*23*). At 37 °C, the CH_4_:C_2_H_6_ ratio was ~110, with an increasing trend towards higher temperatures. As the ROS-driven CH_4_:C_2_H_6_ ratios are substantially lower than those observed for archaeal methanogenesis(*38*), the CH_4_:C_2_H_6_ ratios could serve as indicator to distinguish microbial from abiotic processes.

**Fig. 2.**
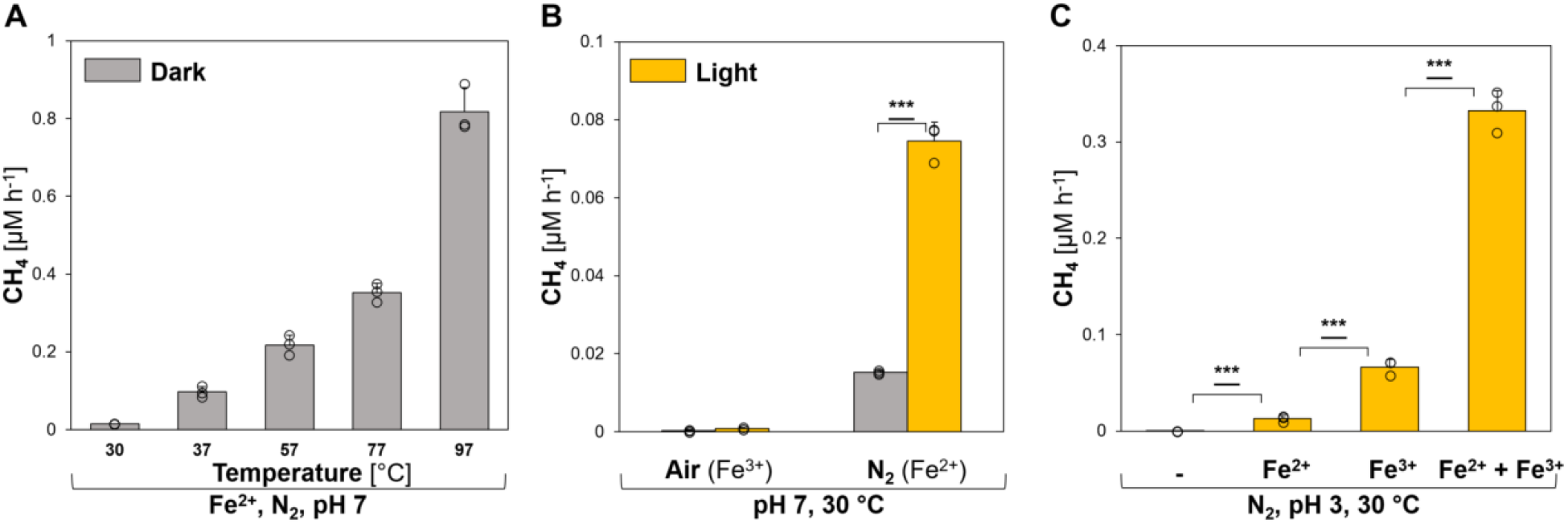
Heat and light drive CH_4_ formation under abiotic conditions. **(A)** Thermolysis: CH_4_ is formed from DMSO under high temperatures **(B)** Water photolysis: The formation of CH_4_ is increased by light **(C)** [Fe(H_2_O)_6_]^3+^ photolysis: Under acidic conditions, light-driven CH_4_ formation is enhanced by [Fe(H_2_O)_6_]^3+^ photochemistry. All experiments were conducted in closed glass vials containing buffered solutions (pH 7 or pH 3) supplemented with DMSO and Fe^2+^ or Fe^3+^ at 30 °C (B, C) under a N_2_ or air atmosphere. Statistical analysis was performed using paired two-tailed t-tests, ***: *p* ≤ 0.001. The bars are the mean + standard deviation of three independent measurements, shown as circles.

Light enhanced the abiotic CH_4_ formation rates (Fig. 2B) by photolysis of water and generation of H_2_O_2_ at 30 °C (Fig. S2). Notably, CH_4_ amounts increased ~4-fold from ~0.02 μM h^-1^ to ~0.08 μM h^-1^ upon broad-spectrum illumination (~350 nm < λ < ~1010 nm at 82 ± 4 μmol photons m^-2^ s^-1^, Fig. S3). This data provides evidence that light-driven CH_4_ formation from methylated S-compounds can occur even in the absence of biomolecules. The addition of oxygen to the samples stopped the formation of CH_4_ in this pH-neutral model system supplemented with Fe^3+^ (Fig. 2B).

In contrast, under acidic (pH 3), illuminated conditions CH_4_ formation rates increased ~5-fold upon Fe^3+^-supplementation in comparison to Fe^2+^-addition, indicating light-driven ROS and Fe^2+^ formation from [Fe(H_2_O)_6_]^3+^ complexes (Fig. 2C)(*35*). Upon supplementation of 1 mM Fe^3+^ and 1 mM Fe^2+^, keeping the overall iron concentration unchanged at 2 mM, CH_4_ formation rates increased to ~0.33 μM h^-1^. This 5-fold rate increase is driven by both ROS-inducing Fe^3+^ and Fenton-driving Fe^2+^. Under pH-neutral conditions, mixing Fe^2+^ and Fe^3+^ only increased CH_4_ formation rates by ~1.3-fold in comparison to Fe^2+^-supplemented samples, while only trace amounts of CH_4_ were obtained from Fe^3+^-supplemented samples (Fig. S4). Thus, illuminated [Fe(H_2_O)_6_]^3+^ complexes generate both Fe^2+^ and ROS, thereby contributing to the ROS-driven CH_4_ formation under acidic conditions.

Taken together, we demonstrated that heat and light drive the formation of CH_4_ and C_2_H_6_ in an anoxic, abiotic environment under ambient temperatures and pressures. These results establish a ROS-driven mechanism based on Fenton chemistry that can occur delocalized from serpentinization across Earth’s humid realm and thereby substantially differs from previously suggested mechanisms that are spatially restricted. Thus, this non-enzymatic hydrocarbon formation mechanism could have released CH_4_ and C_2_H_6_ into the atmosphere of the Hadean and Archean. Besides CH_4_, C_2_H_6_ is considered an important factor in keeping the early Earth warm, since C_2_H_6_ absorbs from 11 to 13 μm in an atmospheric window (roughly 8-13 μm) where H_2_O and CO_2_ do not absorb strongly(*2*). Together, the hydrocarbons produced by these pathways might offer a solution to the “faint young sun paradox”(*3, 4*).

### (Bio)molecules enhance the heat-driven CH_4_ formation

Even before life emerged, several metabolites, e.g. citrate and malate, could have been formed via an ancient, non-enzymatic TCA cycle predecessor driven by ROS(*39, 40*). Catalyzed by iron particles, the formation of pyruvate from CO_2_ was recently reported(*41*). Intriguingly, citrate and malate, as well as other primordial (bio)molecules with a putative prebiotic origin, including ATP(*42*) or serine(*43*), have been reported to act as Fenton-promoting Fe^2+^-chelators(*26*). We therefore investigated if these hydroxylated and carboxylated (bio)molecules enhance the ROS-driven CH_4_ formation rates (Fig. 3A).

**Fig. 3.**
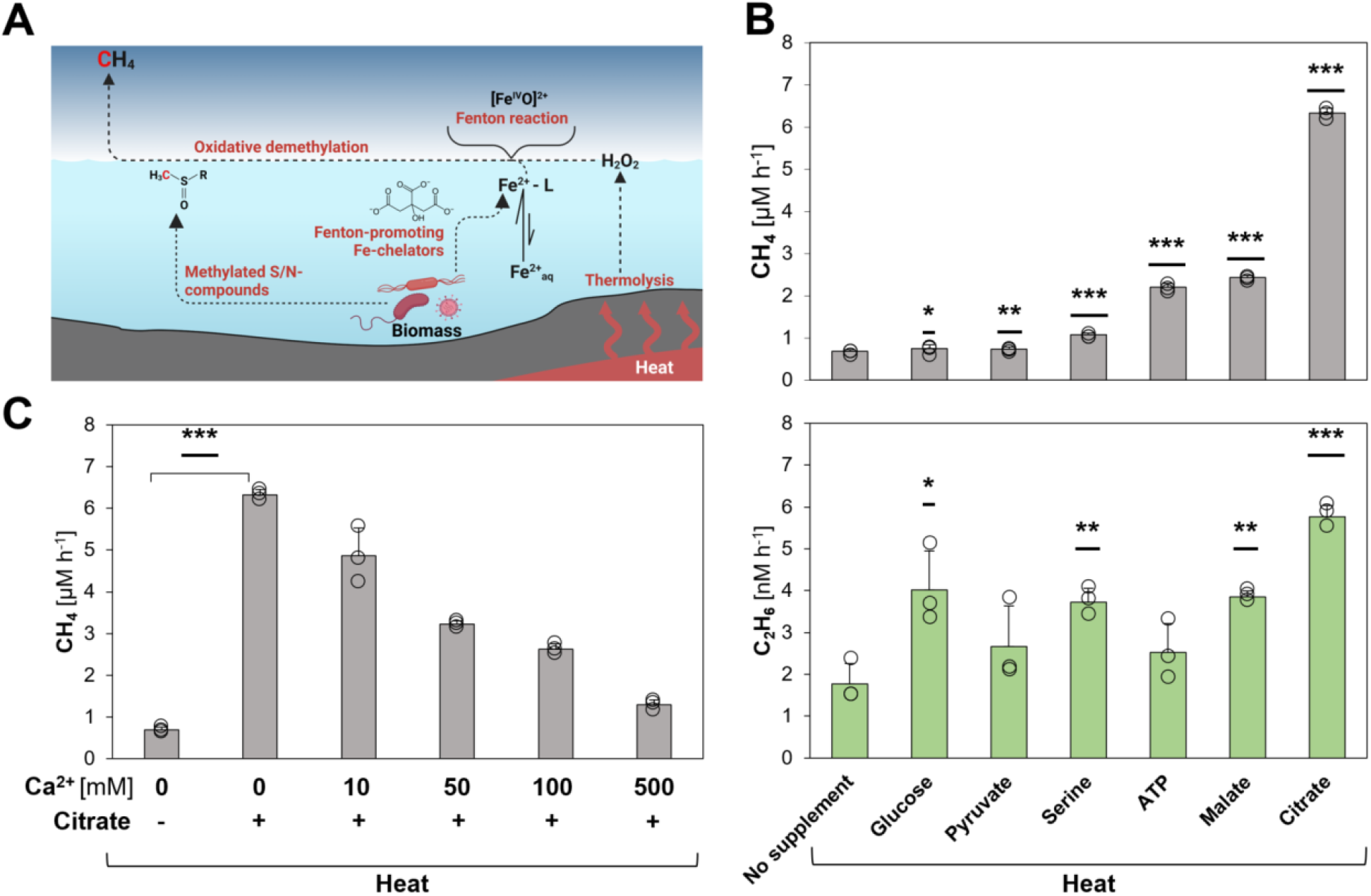
(Bio)molecules enhance heat-driven CH_4_ formation. **(A)** Overview of CH_4_ formation driven by heat. Living organisms produce S-/N-methylated compounds that serve as substrates for CH_4_ formation and Fe^2+^-chelators that promote Fenton chemistry and enhance CH_4_ formation. **(B)** Heat-driven CH_4_ (upper panel) and C_2_H_6_ (lower panel) formation is enhanced upon supplementation with (bio)molecules. **(C)** Citrate enhances heat-driven CH_4_ formation acting as iron-chelator. Upon the addition of Ca^2+^, CH_4_ levels decrease due to the replacement of Fenton-promoting Fe^2+^-citrate complexes with Ca^2+^-citrate complexes. All experiments were conducted in closed glass vials containing a buffered solution (pH 7) supplemented with DMSO, Fe^2+^ and, optionally, citrate and Ca^2+^ under a pure nitrogen atmosphere at 97 °C (heat). Statistical analysis was performed using paired two-tailed t-tests, *: *p* ≤ 0.05, **: *p* ≤ 0.01, ***: *p* ≤ 0.001. The bars are the mean + standard deviation of three independent measurements, shown as circles.

Indeed, the addition of pyruvate, glucose, serine, ATP, malate or citrate to the heat-driven (97 °C) model system (Fig. 3B) increased the abiotic CH_4_ formation rate, e.g. more than 11-fold for citrate. Corresponding C_2_H_6_ rates significantly increased for glucose, serine, malate and citrate, resulting in CH_4_:C_2_H_6_ ratios between ~190 (glucose) and ~1100 (citrate, Fig. 3B). To test if these enhancing effects were indeed driven by Fe^2+^ chelation, we supplemented the assays with the Fe^2+^-competitor Ca^2+^ (Fig. 3C). Since (bio)molecules like citrate can alternatively chelate Ca^2+^ ions, we expected that increasing Ca^2+^ concentrations result in decreasing CH_4_ formation rates by replacing Fenton-promoting Fe^2+^-citrate complexes with Ca^2+^-citrate complexes. Upon addition of 10 mM and 500 mM Ca^2+^, CH_4_ formation rates significantly decreased from ~6.32 μM h^-1^ to ~4.86 μM h^-1^ and ~1.29 μM h^-1^, respectively. Thus, 500 mM Ca^2+^ suppressed ~90 % of the Fenton-promoting effect of citrate supplementation. The Ca^2+^ concentration-dependent decrease of the heat-driven CH_4_ formation rate supports the role of citrate as a Fenton-promoting Fe^2+^-chelator.

Together, ROS generated by heat interact with iron and thereby drive the formation of methyl radicals from S-/N-methylated compounds, resulting in CH_4_ and C_2_H_6_. Moreover, several hydroxylated or carboxylated (bio)molecules with a putative prebiotic origin were shown to act as Fenton-promoting Fe^2+^-chelators, indicating that ROS-driven CH_4_ formation may have already been widespread within the timeframe of the transition from prebiotic chemistry to the origin of life. The rise of life would have fostered the abiotic, non-enzymatic CH_4_ formation due to the consequential formation and release of biomolecules serving as chelators and substrates.

### A light-driven iron redox cycle enhances the Fenton-driven CH_4_ formation

During Fenton chemistry, Fe^2+^ is either oxidized to [Fe^IV^=O]^2+^ or ferric iron (Fe^3+^). As Fe^3+^ cannot drive Fenton reactions (*23, 24*), CH_4_ formation rates decrease with increasing reaction time and increasing concentrations of Fe^3+^. While this effect may have been minor in the ferruginous Archean oceans, Fe^3+^ likely dominated the iron pool in the photic zone of the oceans latest by the rise of photoferrotrophy and was also prevalent in several ecological niches, e.g. volcanic lakes(*34*). The evolution of photosynthesis and the subsequent biological production of O_2_ oxidized the majority of the available Fe^2+^ to Fe^3+^. Thus, abiotic ROS-driven CH_4_ formation would have been hindered in the sunlit realm by the late Archean in the absence of an iron redox cycle at neutral pH. Intriguingly, besides acting as Fenton-promoting Fe^2+^-chelators(*26*), (bio)molecules like citrate were reported to reduce Fe^3+^ to Fe^2+^ via LMCT under oxic and anoxic conditions(*27*). Therefore, (bio)molecules may have facilitated widespread iron redox cycling, e.g. by forming Fe(III)-carboxylate complexes. Furthermore, previous studies showed that, upon illumination of water hydroxyl radicals (·OH) and hydrogen atoms are generated, forming H_2_O_2_ and H2(*30–33*). Thus, we hypothesized that light could drive CH_4_ formation in the absence of Fe^2+^ by simultaneously (i) generating ROS from water and (ii) reducing Fe^3+^ to Fe^2+^ via LMCT, thereby recycling Fe^3+^ and keeping the Fenton reaction running (Fig. 4).

**Fig. 4.**
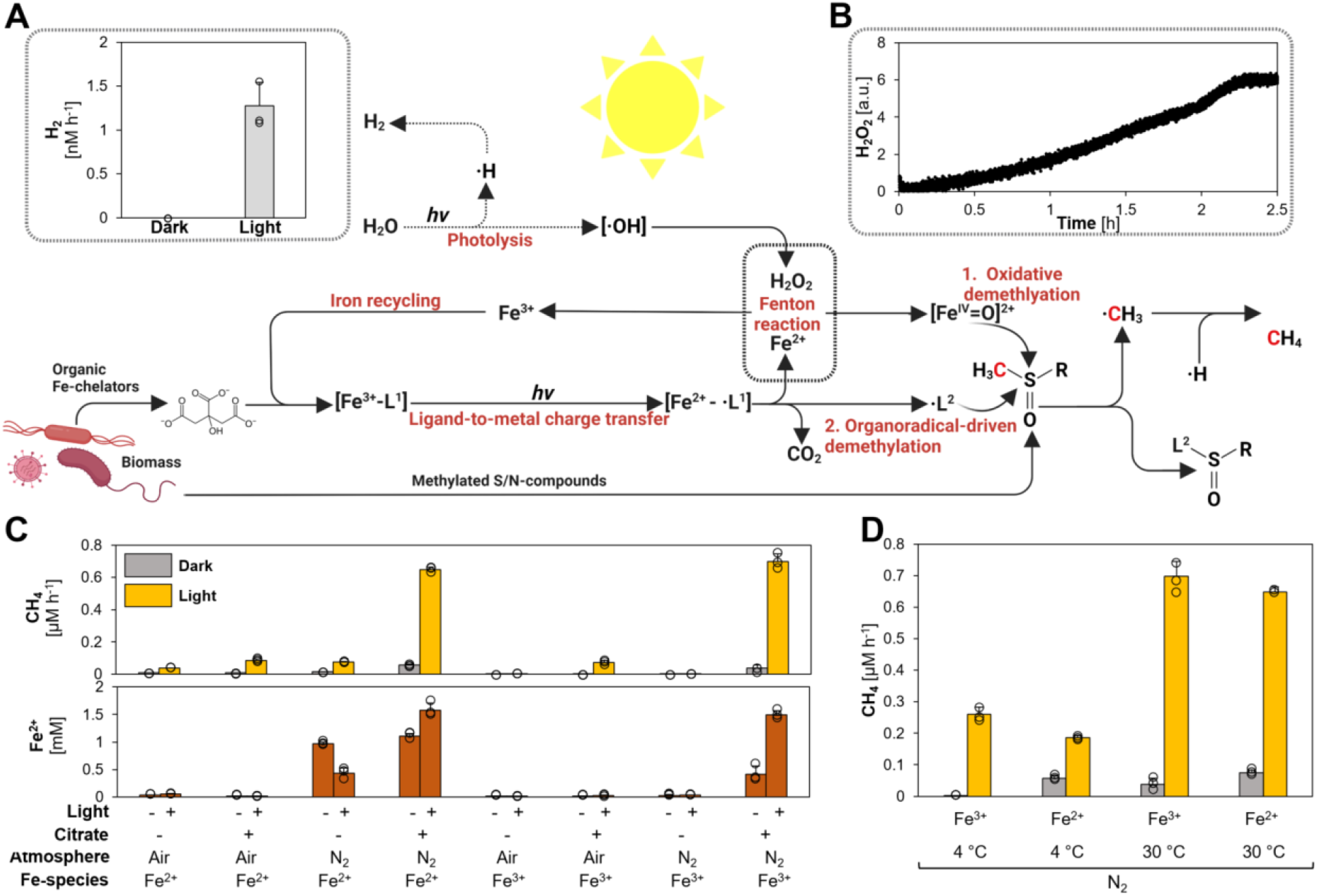
A light-driven iron redox cycle drives and enhances CH_4_ formation. Upon illumination, water is photolytically split into hydroxyl radicals (·OH) and hydrogen forming H_2_ and H_2_O_2_. Organic Fe^3+^-complexes (Fe^3+^-[L^1^]) are converted into Fe^2+^ and organic radicals (·L^1^) via ligand-to-metal charge transfer (LMCT). The generated Fe^2+^ reacts with H_2_O_2_ to ·OH or [Fe^IV^=O]^2+^ and thereby drives the generation of methyl radicals (·CH3) from S-/N-methylated compounds. The LMCT-generated ·L^1^ decomposes into CO_2_ and another organic radical (·L^2^) that additionally facilitates CH_4_ formation upon reacting with S-/N-methylated compounds. Under light, **(A)** H_2_ (grey bars) and (**B**) H_2_O_2_ is formed in pure buffer. **(C**) Upon illumination, CH_4_ formation rates (yellow bars) are increased. Fe^2+^ formation (brown bars) depends on anoxic conditions and is driven by LMCT induced by the addition of citrate. **(D)** Light and heat have synergistic effects on CH_4_ formation. While heat drives CH_4_ formation upon Fe^2+^-supplementation, light increases CH_4_ formation upon Fe^3+^- and Fe^2+^-addition. All experiments were conducted in closed glass vials containing a buffered solution (pH 7) supplemented with DMSO, Fe^2+^ or Fe^3+^, N_2_ or air atmosphere in the presence or absence of citrate incubated under light or in the dark at 4 °C or 30°C. The bars are the mean + standard deviation of three independent measurements, shown as circles.

To verify our hypothesis, we first confirmed light-dependent ROS production in our model system in the absence of substrate, iron and organic ligands by measuring final reaction products of photolysis: H_2_ and H_2_O_2_ (Fig. 4A and 4B). We measured H_2_ production at a rate of ~1.3 nM h^-1^ in anoxic samples under broad-spectrum illumination but not in samples kept in the dark (Fig. 4A). A continuous formation of H_2_O_2_ was measured online using microsensors, which confirmed light-dependent production dynamics in pure buffer (Fig. 4B). Via fluorescence-based H_2_O_2_ endpoint measurements, we found that both iron and DMSO reduced the H_2_O_2_ concentrations. The decrease in H_2_O_2_ levels can be attributed to Fenton reactions between H_2_O_2_, Fe^2+^ and the radical scavenger DMSO (Fig. S5).

Building on this, we closely investigated the interplay of LMCT and iron photochemistry on CH_4_ formation. For this purpose, we analyzed our chemical model system containing a buffered solution (pH 7), Fe^2+^ or Fe^3+^, DMSO, in the presence or absence of citrate for the formation of CH_4_ and the concentration of available Fe^2+^ (Fig. 4C). The influence of the following parameters on the formation of CH_4_ was tested: (i) O_2_ (~21% in air), (ii) oxidation state of the supplemented iron species (Fe^2+^ *vs.* Fe^3^), (iii) light and (iv) presence/absence of citrate. (i) CH_4_ formation rates under anoxic conditions always exceeded rates under oxic conditions. (ii) Without citrate, initial Fe^2+^-supplementation was required to form significant CH_4_ levels. (iii) CH_4_ formation always increased with light. (iv) Upon citrate addition, CH_4_ formation was enhanced in illuminated and anoxic samples containing DMSO and Fe^2+^ or Fe^3+^. Besides elevated CH_4_ formation rates, citrate addition also increased the final Fe^2+^ concentrations, e.g. from ~0 mM Fe^2+^ to ~1.5 mM Fe^2+^ in illuminated and anoxic samples.

After determining the influence of the four parameters (i) O_2_, (ii) iron (iii) light and (iv) (bio)molecules, we further investigated them to gain a better understanding of their contribution and role in the light-driven CH_4_ formation.

i. O_2_: The influence of O_2_ on LMCT and CH_4_ formation was studied in citrate-supplemented samples by adding various amounts of air. Fe^2+^ concentrations and CH_4_ formation rates decreased with increasing O_2_ levels (Fig. S6). In comparison to 0 % O_2_, the Fe^2+^ concentration dropped drastically already at 0.2 % O_2_ and was ~96 % lower at 2 % O_2_, while CH_4_ formation rates decreased approximately linearly with the O_2_ level. This indicates the presence of a Fe-cycle, in which most LMCT-formed Fe^2+^ is instantly re-oxidized, either by O_2_ or Fenton reactions. The balance between these Fe^2+^ sinks depend on O_2_ availability and governs CH_4_ formation rates. In the presence of O_2_, we also detected methanol (CH_3_OH) formation rates ranging from ~0.003 μM h^-1^ (0.2 % O_2_) to ~0.07 μM h^-1^ (21 % O_2_). CH3OH is preferentially formed through the reaction of ·CH_3_ with O_2_(*23, 44*). Without the addition of O_2_, no CH3OH was detected, indicating anoxic conditions in our standard assays.
ii. Iron: The role of the LMCT-rate and the corresponding Fe^2+^ availability for CH_4_ formation was tested by supplementing the assays with various Fe^3+^ concentrations (Fig. S7). At lower Fe^3+^ concentrations, CH_4_ formation rates increased steeper than the measured Fe^2+^ concentrations. At high Fe^3+^ concentrations, CH_4_ formation rates leveled off, while Fe^2+^ concentrations continued to increase. This indicates that Fe^2+^ is limiting the demethylation rates at low iron concentrations, because it is immediately re-oxidized. While light-dependent ROS production is limiting CH_4_ formation at high iron concentrations. Most importantly, these data highlight that a light- and ROS-driven iron cycle can facilitate high rates of CH_4_ formation, even in the presence of O_2_ and the absence of detectable Fe^2+^, which opens the possibility of widespread abiotic CH_4_ production after the great oxidation event as well as in diverse modern habitats. Besides iron, several other transition metals were reported to drive Fenton chemistry(*45, 46*), resulting in the release of CH_4_. Thus, we test different transition metals in our chemical model system, containing DMSO as substrate and ascorbate as a strong metal reductant(*47, 48*). We observed that copper, cobalt and cerium also enhanced CH_4_ formation rates (Fig. S8). However, the effect was less pronounced compared to iron. The high activity of iron combined with its ubiquitous abundance in the Precambrian highlights the global distribution and importance of this mechanism.
iii. Light: It is established the light quality has an important influence on photolysis. Short wavelength light in the ultraviolet spectrum was reported to drive water photolysis and LMCT more efficiently than longer wavelengths(*49*). We expected that shorter wavelength light would increase both CH_4_ formation rates and Fe^2+^ levels. Indeed, CH_4_ formation rates surged from ~0.3 μM h^-1^ (λ_max_ = 534 nm) to ~1.23 μM h^-1^ (λ_max_ = 388 nm, Fig. S9) and Fe^2+^ concentrations almost tripled from ~1.3 mM (λ_max_ = 534 nm) to ~4.2 mM (λ_max_ = 388 nm, Fig. S9). Although the broad-spectrum light had a 1.5-fold higher energy flux (57 ± 2 kJ m^-2^ h^-1^) compared to the 388 nm-LED light (37 ± 2 kJ m^-2^ h^-1^), the CH_4_ formation rate under the broad-spectrum light was only half (0.7 μM h^-1^). Given that the stratospheric ozone layer was absent during the Hadean and Archaean, higher fluxes of short wavelength light (*i.e.* ultraviolet light), reached aqueous environments and may have further enhanced the ROS-driven CH_4_ formation.
iv. (Bio)molecules: After illumination of Fe^3+^-ligand complexes, one electron is transferred via LMCT from a carboxylated ligand (L^1^) to Fe^3+^, an organic radical (·L^1^), *i.e.* citrate radical, is generated. As described in the literature (*28*), we observed the subsequent CO_2_ disassembly from citrate radicals (Fig. S10). We speculated that the remaining organic radical (·L^2^) could react with DMSO, resulting in ·CH3 and the formation of CH_4_ (Fig. 4). Since we cannot directly detect organic radicals, we mimicked the proposed reaction in an anoxic model system only containing DMSO and the radical generating 2,2’-Azobis(2-amidinopropane) dihydrochloride (APPH) that readily decomposes into carbon-centered organic radicals at 40 °C (Fig. S11). Indeed, we observed CH_4_ formation in a mixture of DMSO and APPH, while only trace amounts of CH_4_ were observed from either DMSO or AAPH alone, this supports an organic radical-driven CH_4_ formation mechanism. In short, carboxylic acids like citrate facilitate LMCT to reduce Fe^3+^ to Fe^2+^ and form organic radicals, both compounds drive CH_4_ formation. Overall, CH_4_ can be formed under anoxic conditions via (i) water thermolysis, (ii) water photolysis, (iii) [Fe(H_2_O)_6_]^3+^ photolysis and (iv) LMCT-induced carbon-centered radicals. Apart from serving as chelators, some (bio)molecules could also serve as substrates for Fenton reactions. Thus, we investigated four S-/N-methylated compounds in the presence of the chelator citrate. Upon illumination, CH_4_ was formed from dimethyl sulfide, methionine, 2-methylthioethanol and trimethylamine (Fig. S12). These observations indicate that ROS-driven CH_4_ formation significantly increased after the origin of life by providing biomolecules as chelators and substrates.

Finally, synergistic effects between light and heat were observed (Fig. 4D). For Fe^2+^-supplemented samples, CH_4_ rates at 4 °C increased from ~0.056 μM h^-1^ in the dark over ~0.19 μM h^-1^ under light to ~0.65 μM h^-1^ in illuminated samples at 30 °C. For Fe^3+^-supplemented samples, only CH_4_ rates below 0.03 μM h^-1^ were obtained in the dark, while CH_4_ formation rates were slightly above Fe^2+^-supplemented samples in the light, again demonstrating the effects of LMCT and LMCT-induced carbon-centered radicals. Thus, the two factors heat and light synergistically combine for a stable and enhanced ROS and CH_4_ formation.

### ROS-generated CH_4_ is derived from biomass with an abiotic isotope fractionation

Considering the impact of (bio)molecules on the LMCT-driven Fenton reaction, organic radical generation and the role of biomolecules as substrates, we expect the discussed mechanisms to have played and still play the most important role in the vicinity of decaying biomass. To demonstrate that CH_4_ is indeed formed from dead biomass in the presence of a variety of biomolecules and not just in our well-defined model systems, we conducted deuterium labeling experiments. For this purpose, we grew the bacterium *B. subtilis* in *Luria-Bertani* medium supplemented with 10% D_2_O and inactivated the cells by sonication and freezing (see Methods).

The obtained dead biomass was supplemented with Fe^3+^ and ascorbate and incubated under broad-spectrum light. Around 40 fmol CH_4_ h^-1^ mg^-1^ dry weight was obtained from labeled and unlabeled biomass (Fig. 5A). In addition, stable hydrogen isotope values (*δ^2^H*) of CH_4_ from D_2_O-treated biomass showed strong enrichment in deuterium (~5900 ‰) in comparison to unlabeled biomass (~-225 ‰), demonstrating a direct conversion of isotopically labeled biomass to CH_4_. This suggests that the availability of biomass, upon the emergence of life, has increased the CH_4_ formation by delivering both (i) S-/N-methylated compounds and (ii) Fenton-promoting iron chelators. The presence of CH_4_ has been suggested to be crucial for the evolution of life, since it could serve as life’s first carbon source via methanotrophy(*50–52*). Following this line of thought, we could demonstrate that methanotrophic *Methylocystis hirsuta* grew on CH_4_ generated by our light-driven model system, transferred to the headspace of the *M. hirsuta* culture (Fig. S13). In fact, the “last methane-metabolizing ancestor” had likely the genes to perform methanogenesis and anaerobic methane oxidation(*53*), suggesting that, under high CH_4_ concentrations, methanotrophy could have emerged prior to methanogenesis.

**Fig. 5.**
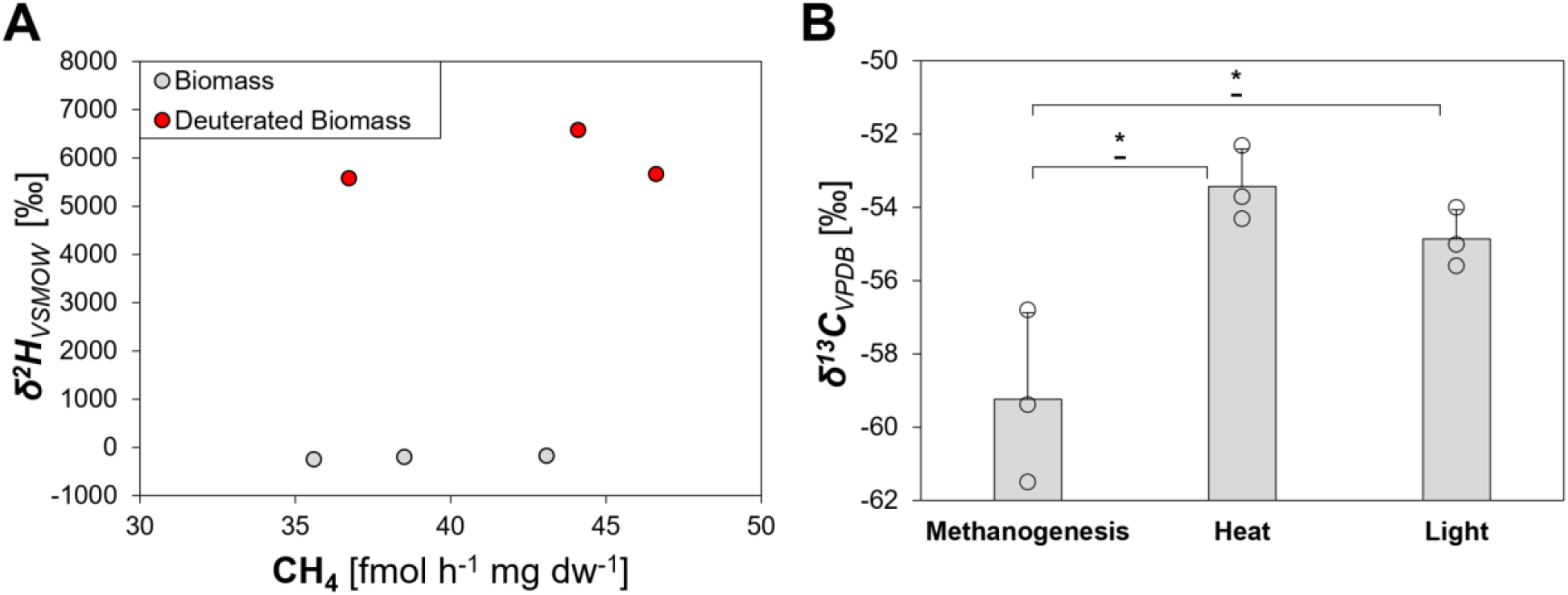
Isotope labeling studies confirm dead biomass as substrate and show an abiotic isotope fractionation for ROS-driven CH_4_ formation. **(A)** Unlabeled or deuterium-enriched CH_4_ is formed from unlabeled biomass (grey dots) or deuterated biomass (red dots), respectively. **(B)** Stable carbon isotope values of cultures from the methanogen *Methanothermobacter marburgensis,* heat-, or light-generated CH_4_. All experiments were conducted in closed glass vials containing a buffered solution (A, B – heat, light) or culture medium (B – methanogenesis), supplemented with Fe^3+^ and ascorbate (A) or Fe^2+^ and citrate (B – heat, light) under a nitrogen atmosphere, incubated under light at 30 °C or in the dark at 97 °C. Statistical analysis was performed using paired two-tailed t-tests, *: *p* ≤ 0.05. The bars are the mean + standard deviation of three independent measurements, shown as circles.

Finally, we speculated that ROS-driven CH_4_ formation leads to different stable carbon isotope values (*δ^13^C*) compared to biological processes, *i.e.,* methanogenesis. The observed *δ^13^C* values for CH_4_ generated by heat or light were less negative (~-54 ± 1.1 compared to the *δ^13^C* value of the methanogen *Methanothermobacter marburgensis* (~-59.2 ± 2.3 Fig. 5B). While the isotopic fractionation during abiotic ROS-driven CH_4_ formation remains to be studied in depth, these results suggest a lower carbon isotope fractionation for ROS-driven CH_4_ formation than for enzymatic methanogenesis. Together with the observed CH_4_:C_2_H_6_ ratios, isotopic signatures may therefore serve to differentiate between CH_4_ formed enzymatically or abiotically on Earth and extraterrestrial planets.

## Conclusions

The findings of this study have significant implications for our understanding of the origin of CH_4_ and C_2_H_6_ on Earth and other planets. We demonstrated that the interplay of Fe^2+^ and H_2_O_2_, generated by heat and light, drives CH_4_ formation from methylated S-/N-compounds via Fenton chemistry under conditions that were globally prevalent in the Hadean and Archean and, principally, remain until today. These new pathways allow a CH_4_ and C_2_H_6_ production in many aqueous environments including oceans, lakes, rivers, and ponds, delocalized from restricted hotspots for (bio)molecule formation such as hydrothermal vents or ultramafic rocks. After the emergence of life, this phenomenon would have greatly intensified in the anoxic Archaean and the subsequent “boring billion” (*54, 55*). The increasing amounts of biomass provided methylated S-/N-substrates, Fe-chelating biomolecules reducing Fe^3+^ to Fe^2+^ and releasing organic radicals and thus enhance ROS-driven CH_4_ formation. Possibly, these reactions facilitated elevated CH_4_ and C_2_H_6_ levels during the Hadean and Archean. These hydrocarbons would have contributed to atmospheric temperatures on Earth and allowed the evolution of life in a liquid hydrosphere, potentially influencing the evolution of metabolism by allowing the rise of methanotrophy prior to methanogenesis. This work lays the foundation to explore further the mechanism’s role in shaping the evolution of the atmosphere on Earth and other planets and its influence on the current climate change.

## Acknowledgments

We thank N. Oehlmann, F. Schmidt, H. Addison, A. Lago Maciel, G. Marijan, A. Goldman, K. Guo, F. Arriaza-Gallardo, M. Schneider, C. Hoyer, M. Schroll and M. Greule for practical and theoretical support; I. Bischofs, S. Shima and W. Liesack for providing strains; G. Hochberg, M. Preiner and T. Erb for providing critical comments and feedback. Figures 1, 3a and 4 were created with BioRender.com.

## Funding

German Research Foundation grant 446841743 (JGR, LE)

German Research Foundation grant KE884 19-1 (FK)

The Max Planck Society (LE, JMK, JGR)

Friedrich Naumann Foundation (LE)

## Author contributions

Conceptualization: LE, JGR

Methodology: LE, UB, JH, TD, JMK, FK, JGR

Investigation: LE, UB (Figs. S2/5/10/11), JMK (Fig. 4B, S3, S9), JH (Fig. S6 – methanol)

Visualization: LE, JGR

Funding acquisition: LE, TD, JMK, FK, JGR

Project administration: JGR

Supervision: JGR

Writing – original draft: LE, JGR

Writing – review & editing: LE, JMK, FK, JGR

## Data and materials availability

All data are available in the main text or the supplementary materials and on request from the corresponding authors.

## Supplementary Materials

Materials and Methods

Figs. S1 to S13

References (*56 - 60*)

