## Supplemental Material for "Methane formation driven by light and heat prior to the origin of life"

**The PDF file includes:**

Materials and Methods
Figs. S1 to S13

### Materials and Methods

#### General assay conditions

Unless otherwise indicated, 4 mL samples were incubated in closed 20 mL glass vials at 30 °C under a pure nitrogen ($N_2$) atmosphere and subsequently analyzed via gas chromatography (GC).

#### Heat assays

500 mM DMSO and 10 mM $FeSO_4$ were added to 20 mM degassed potassium phosphate buffer (pH 7) in an anaerobic tent. The headspace of the closed vials was then cycled three times with vacuum and $N_2$. Samples were incubated at 37 °C, 57 °C, 77 °C and 97 °C for 6 h in an incubator in the dark. Optionally, 20 mM citrate, malate, ATP, serine, glucose or pyruvate were also supplemented. $Ca^{2+}$ was added in the form of $CaCl_2$. Samples were measured within the linear range of $CH_4$ formation rate via gas chromatography.

#### Light assays

500 mM DMSO, 2 mM of either $FeCl_3$ or $FeSO_4$ and, optionally, 10 mM citrate were added to 20 mM degassed potassium phosphate buffer (pH 7). Anoxic conditions were generated by drawing vacuum eight times for 1 min and a subsequent filling with $N_2$. For experiments investigating $[Fe(H_2O)_6]^{3+}$ complexes, samples were incubated under anoxic, acidic conditions (pH 3) and supplemented with 500 mM DMSO and either 2 mM $FeCl_3$, 2 mM $FeSO_4$ or 1 mM $FeCl_3$ and 1 mM $FeSO_4$, each. Samples were incubated under air or $N_2$ in the dark or under constant broad-spectrum illumination from light bulbs (Osram, Superlux, Super E SIL 60; $\Phi$ = 82 ± 4 µmol photons $m^{-2}$ $s^{-1}$, $H$ = 52 ± 2 kJ $m^{-2}$ $h^{-1}$; Fig. S3) for 1 day. Samples were measured within the linear range of the $CH_4$ formation via gas chromatography. Specific wavelengths were provided by diodes (H2A1 series, Roithner Lasertechnik, Austria) emitting UV-A, blue, cyan, green, red or near-infrared light ($\lambda_{max}$ = 388 nm, $\Phi$ = 35 ± 1 µmol photons $m^{-2}$ $s^{-1}$, $H$ = 36 ± 2 kJ $m^{-2}$ $h^{-1}$; $\lambda_{max}$ = 436 nm, $\Phi$ = 45 ± 1 µmol photons $m^{-2}$ $s^{-1}$, $H$ = 45 ± 1 kJ $m^{-2}$ $h^{-1}$; $\lambda_{max}$ = 500 nm, $\Phi$ = 64 ± 4 µmol photons $m^{-2}$ $s^{-1}$, $H$ = 55 ± 3 kJ $m^{-2}$ $h^{-1}$; $\lambda_{max}$ = 534 nm, $\Phi$ = 63 ± 1 µmol photons $m^{-2}$ $s^{-1}$, $H$ = 50 ± 1 kJ $m^{-2}$ $h^{-1}$; $\lambda_{max}$ = 675 nm, $\Phi$ = 45 ± 4 µmol photons $m^{-2}$ $s^{-1}$, $H$ = 29 ± 3 kJ $m^{-2}$ $h^{-1}$; or $\lambda_{max}$ = 868 nm, $\Phi$ = 69 ± 7 µmol photons $m^{-2}$ $s^{-1}$, $H$ = 35 ± 3 kJ $m^{-2}$ $h^{-1}$) Light intensity was determined using a fiber optic scalar irradiance microsensor*(56)* connected to a spectrometer (USB4000; Ocean Optics, USA) placed in the center of the incubation vials and calibrated using a spherical light probe (Walz) connected to a LI-250A light meter (Li-Cor Biosciences GmbH, Germany)*(57)*. Concentration of $Fe^{2+}$ was quantified with the colorimetric ferrozine method*(58)*.

#### *Bacillus subtilis* biomass assays

*B. subtilis* was grown in 500 mL LB media, supplemented with 10 % $H_2O$ or $D_2O$, grown for 36 h at 37 °C and 180 rpm. The obtained culture was collected by three cycles of centrifugation (10 min, 4000 rpm) and resuspended in 35 mL 20 mM potassium phosphate buffer (pH 7) in order to remove the excess $D_2O$. Biomass was then generated by sonication (4-times, 1 min) and freezing of the samples. Subsequently, 80 mL buffer was supplemented with 10 mL biomass, 20 mM $FeCl_3$ and 50 mM ascorbic acid, saturated with $N_2$ for 30 min and incubated in 100 mL closed glass vials under $N_2$ and constant broad-spectrum illumination for 3 days. The gas headspace was extracted with a syringe and analyzed with regard to $CH_4$ content and $\delta^2 H$ values.

**_Methylocystis hirsuta_ and _Methanothermobacter marburgensis_ cultivation**

*M. hirsuta* growth media contained 0.5 g $Na_2HPO_4 \cdot 2H_2O$, 0.22 g $KH_2PO_4$, 1 g $KNO_3$, 0.4 mg $CaCl_2 \cdot 2H_2O$, 2 mg $MgSO_4 \cdot 7H_2O$ per liter, supplemented with 5 mg $Na_2EDTA$, 0.06 mg $CuCl_2 \cdot 5H_2O$, 2 mg $FeSO_4 \cdot 7H_2O$, 0.1 mg $ZnSO_4 \cdot 7H_2O$, 0.03 mg $MnCl_4 \cdot 4H_2O$, 0.05 mg $H_3BO_3$, 0.2 mg $CoCl_2 \cdot 6H_2O$, 0.02 mg $NiCl_2 \cdot 6H_2O$ and 0.03 mg $Na_2MoO_4 \cdot 2H_2O$ per liter. *M. hirsuta* was cultivated in 100 mL closed glass vials containing 30 mL culture and was incubated at 25 °C and 150 rpm under an air atmosphere. Methane was produced by supplementing 2 L degassed 20 mM potassium phosphate buffer with 1 M DMSO, 25 mM $FeSO_4$ and 50 mM ascorbic acid, incubating the solution under constant illumination in 1 L flasks and collecting the formed $CH_4$ with syringes. *M. hirsuta* cultures were either supplemented with 25 mL light-generated $CH_4$ or 25 mL pure $N_2$. *M. marburgensis* was cultivated as previously described*(59)*.

**Continuous $H_2O_2$ measurements using microsensors**

To visualize $H_2O_2$ production in the illuminated anoxic model system, an $H_2O_2$ microsensor was positioned in the solution. The $H_2O_2$ microsensors were built, calibrated and used as described previously*(60)*. We sealed the vial opening with self-adhesive tape, rigorously bubbled the liquid with $N_2$ and then adjusted a gentle flow of $N_2$ through the headspace to minimize oxygen input from the atmosphere. Light was provided from halogen lamps (KL2500, Schott) at an intensity of 1027 µmol photons $m^{-2}$ $s^{-1}$. We did not attempt to calculate light-dependent $H_2O_2$ production rates due to the open design of the system, which allowed for the exchange of $H_2O_2$ with the headspace across the water interface.

**End-point $H_2O_2$ measurements**

After illumination, 290 µL sample was mixed anaerobically with 9 µL Amplex Ultrared (Thermofisher, A36006, 30 µM final concentration) and 1 µL recombinant APEX2 (0.23 µM final concentration). Fluorescence was then measured with a plate reader (BMG ClarioStar™) at 568 nm excitation / 581 nm emission. A calibration curve was established with $H_2O_2$ following the same procedure. To prevent $O_2$-driven $H_2O_2$ generation while sample preparation, all buffers were saturated with $N_2$ and the plate reader was kept at a partial oxygen pressure of 0.1% with an atmospheric control unit (Clariostar, BMG). Before sample preparation, all sample components (20 mM potassium phosphate buffer, DMSO, 1 M citrate and 100 mM $FeCl_3$) were degassed and kept in an anoxic tent overnight.

**Quantification of $CH_4$, $C_2H_6$, $CO_2$, and $H_2$ (GC-FID)**

Amounts of formed $CH_4$, $C_2H_6$, $CO_2$ and $H_2$ were determined via headspace analysis using a PerkinElmer® Clarus®690 GC system (GC–FID/TCD) with a custom-made column circuit (ARNL6743). The headspace samples were injected by a TurboMatrixX110 (PerkinElmer Inc, Waltham, USA) autosampler, heating the samples to 45 °C for 15 min prior to injection. The samples were then separated on a HayeSep column (7' HayeSep N 1/8'' Sf; PerkinElmer®), followed by molecular sieve (9' Molecular Sieve 13x 1/8'' Sf; PerkinElmer®) kept at 60 °C. Subsequently, the gases were detected with a flame ionization detector (FID, at 250 °C) and a thermal conductivity detector (TCD, at 200 °C). The quantification of $CH_4$, $C_2H_6$, $CO_2$ and $H_2$ was based on linear standard curves that were derived from measuring varying amounts of these gases.

**$CH_3OH$ measurements (GC-FID)**

$CH_3OH$ was quantified with a GC-FID (Shimadzu GC-2010 Plus, FID-2010 Plus, 280°C) containing an AOC 20i autosampler and a ZB-WAXplus (Zebron) column (30 m x ⌀=0.25 mm, df, 0.25µm). A $H_2O$ sample (1 µL) was injected in the split liner (250°C, split 5,15,50). The temperature program was kept at 35°C for 5 min and then increased by 50°C $min^{-1}$ until 200°C which was kept for 3 min. Helium served as carrier gas (flow rate: 1.95 ml $min^{-1}$) and the FID was operated with 400 ml $min^{-1}$ synthetic air, 40 ml $min^{-1}$ $H_2$ and 30 ml $min^{-1}$ $N_2$, serving as a makeup gas. For Split 5, a calibration curve ($R^2$= 0.9931) was generated by diluting $CH_3OH$ (99.9% purity), while an $R^2$=0.9981 for split 15 and an $R^2$=0.9997 for split 50 was determined.

**$\delta\ ^{13}C$ stable isotope measurements (GC-C-IRMS)**

$\delta^{13}C$ values of $CH_4$ were determined by gas chromatography-combustion-isotope ratio mass spectrometry (GC-C-IRMS). Aliquots of headspace gas were transferred to an evacuated sample loop (40 mL) and a cryogenic pre-concentration unit to trap $CH_4$. $CH_4$ was trapped on HayeSep D, separated from interfering compounds by GC and transferred to the GC-C-IRMS. The system consists of a cryogenic pre-concentration unit directly connected to an HP 6890N GC (He flow rate: 1.8 mL $min^{-1}$; Agilent Technologies, Santa Clara, USA) fitted with a GS-Carbonplot capillary column (30 m * 0.32 mm i.d., $d_f$ 1.5 µm; Agilent Technologies) and a PoraPlot capillary column (25 m * 0.25 mm (i.d.), $d_f$ 8 µm; Varian, Lake Forest, USA). The GC flow was coupled using a press-fit connector to a combustion reactor comprised of an oxidation reactor (ceramic tube ($Al_2O_3$), length 320 mm, inner diameter 0.5 mm, with oxygen-activated Cu/Ni/Pt wires inside; reactor temperature 960°C) and a GC Combustion III Interface (ThermoQuest Finnigan) to decompose $CH_4$ into $CO_2$. $^{13}C/^{12}C$ ratios were determined with a Delta$^{PLUS}$XL mass spectrometer (ThermoQuest Finnigan, Bremen, Germany). High-purity $CO_2$ (Messer Griesheim, Frankfurt, Germany) was used as the working monitoring gas. $^{13}C/^{12}C$ ratios ($\delta^{13}C$ values) are expressed in the conventional δ notation in per mil versus VPDB, calculated as:

$$\delta^{13}C_{VPDB} = \left( \frac{\left(\frac{^{13}C}{^{12}C}\right)_{Sample}}{\left(\frac{^{13}C}{^{12}C}\right)_{Standard}} \right) - 1$$

$\delta^{13}C$ values were corrected using three reference standards of high-purity $CH_4$ with $\delta^{13}C$ values of –54.5 ± 0.2 ‰ (Isometric Instruments, Victoria, Canada), –66.5 ± 0.2 ‰ (Isometric Instruments) and –42.3 ± 0.2 ‰ (in-house), calibrated against International Atomic Energy Agency and NIST reference substances.

**$\delta^2H$ stable isotope measurements (GC-TC-IRMS)**

$\delta^2H$ values for $CH_4$ were determined using GC-temperature conversion-isotope ratio mass spectrometry (GC-TC-IRMS). The analytical set-up was the same as the one used for $\delta^{13}C$ stable isotope measurements except that the He flow rate was changed to 0.6 ml $min^{-1}$ and, instead of combustion to $CO_2$ and $H_2O$, $CH_4$ was thermolytically converted (at 1450 °C) to hydrogen and carbon. After IRMS measurements, the obtained $\delta^2H$ values were corrected by using two reference standards of high-purity $CH_4$ with $\delta^2H$ values of –149.9‰ ± 0.2‰ (T-iso2, Isometric Instruments) and –190.6‰ ± 0.2‰ (in house). All $\delta^2H$ values are expressed in the conventional δ notation in per mil versus Vienna Standard Mean Ocean Water (VSMOW), calculated as

$$\delta\, ^2H_{VSMOW} = \left( \frac{\left(\frac{^2H}{^1H}\right)_{Sample}}{\left(\frac{^2H}{^1H}\right)_{Standard}} \right) - 1$$

**Statistics**

Unless indicated otherwise, all experiments were performed with N = 3 replicates (3 biological replicates). To test for significant differences in $CH_4$ formation between two samples, single-factor analysis (students t-test) of variance (ANOVA) was used.

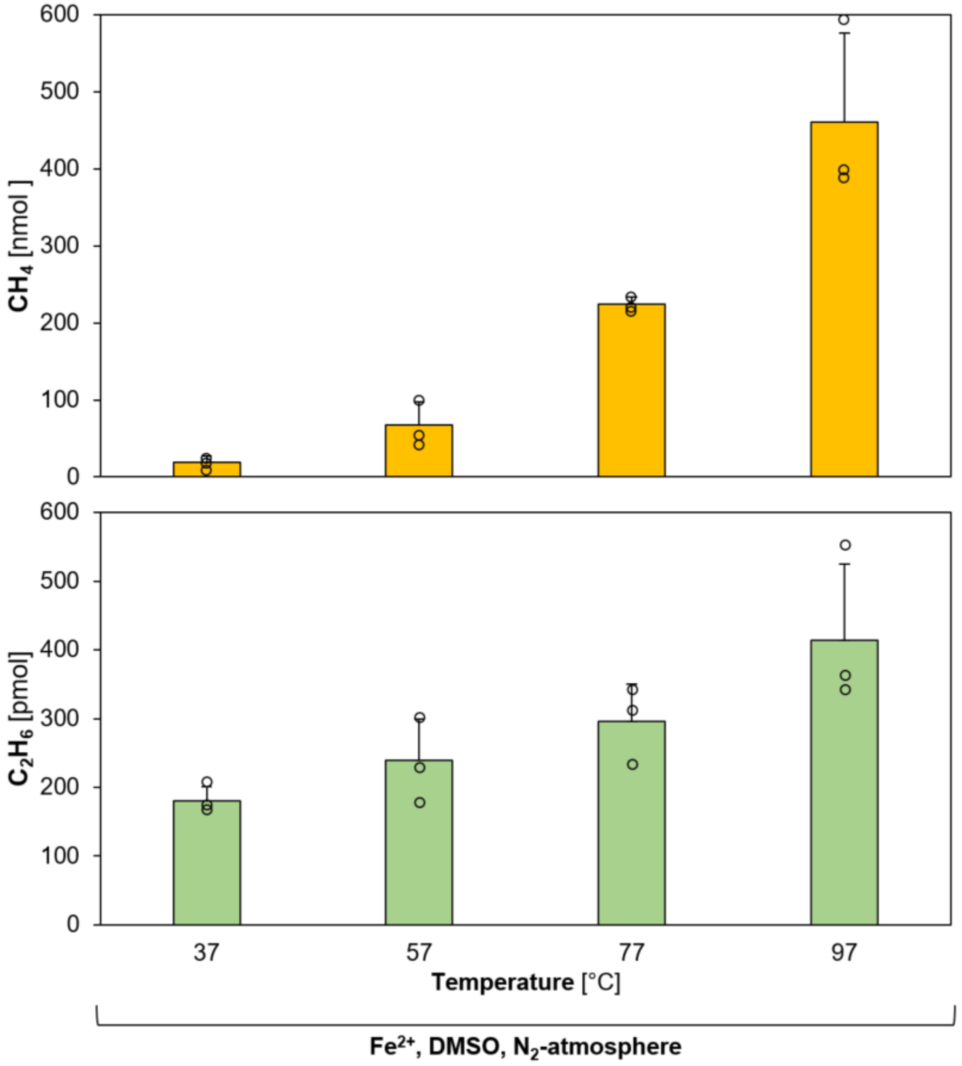


**Fig. S1. Heat-driven $CH_4$ and $C_2H_6$ formation from DMSO.** Formed $CH_4$ amounts increase from ~20 nmol (37 °C) to ~460 nmol (97 °C). Formed $C_2H_6$ amounts increase from ~190 pmol (37 °C) to ~415 pmol (97 °C), thereby resulting in $CH_4$:$C_2H_6$ ratios from ~110 (37 °C) to ~1100 (97 °C). In a total volume of 4 mL, samples containing a 20 mM potassium phosphate buffer (pH 7), 1 M DMSO, 10 mM $FeSO_4$ were incubated at different temperatures in sealed 20 mL glass vials under $N_2$ for 4 days. The bars are the mean + standard deviation of three independent measurements, shown as circles.

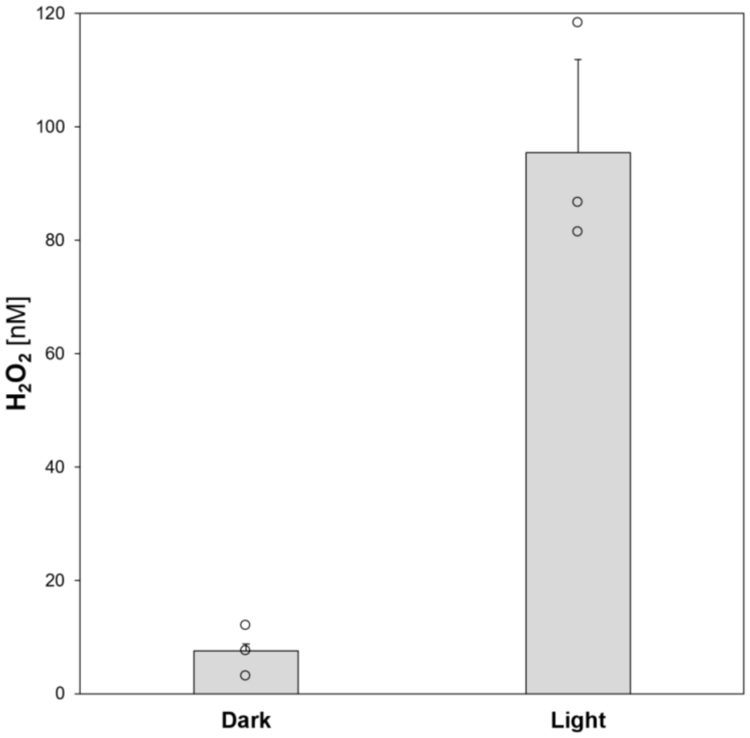


**Fig. S2. Light-driven $H_2O_2$ formation in pure buffer.** Formed $H_2O_2$ concentrations increase from ~7 nM in the dark to ~95 nM under light. In a total volume of 4 mL, samples contained 20 mM potassium phosphate buffer (pH 7) that was bubbled with $N_2$ for 1 h and kept in an anoxic tent for 5 days. Samples were incubated in the dark or under illumination (Fig. S3) for 20 h and analysed via fluorescence-based $H_2O_2$ endpoint measurements (see Methods). The bars are the mean + standard deviation of three independent measurements, shown as circles.

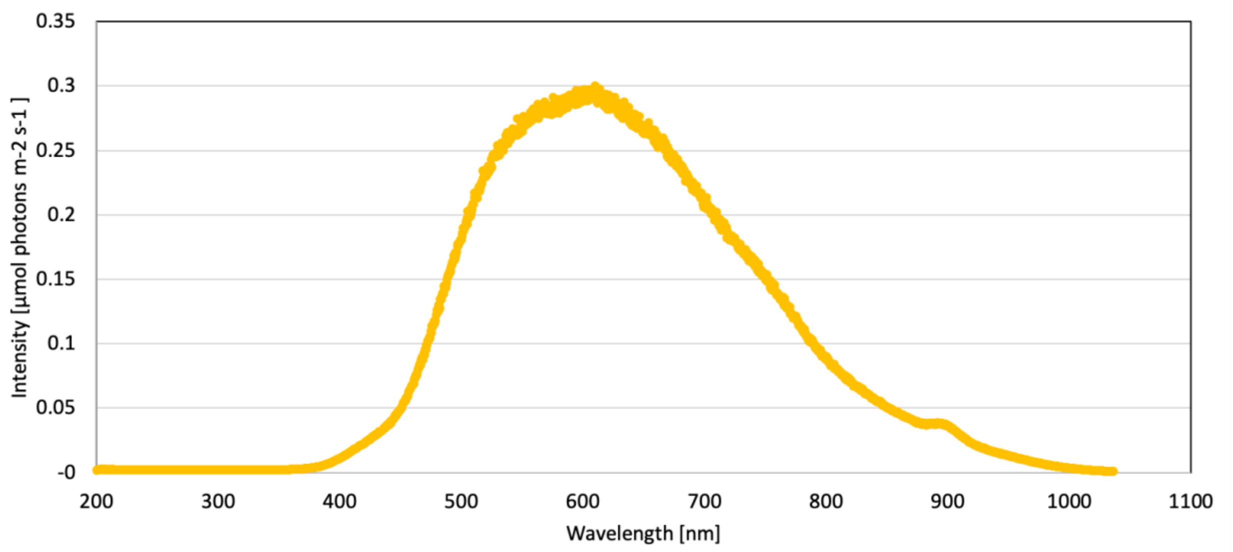


**Fig. S3. Light spectrum of the used light bulbs.** For broad-spectrum sample illumination, Osram (Superlux, Super E SIL 60) light bulbs were used, with an intensity of 82 ± 4 µmol photons $m^{-2}$ $s^{-1}$, and an energy flux of 52 ± 2 kJ $m^{-2}$ $h^{-1}$. The light spectrum was determined with a spectrometer (see Methods).

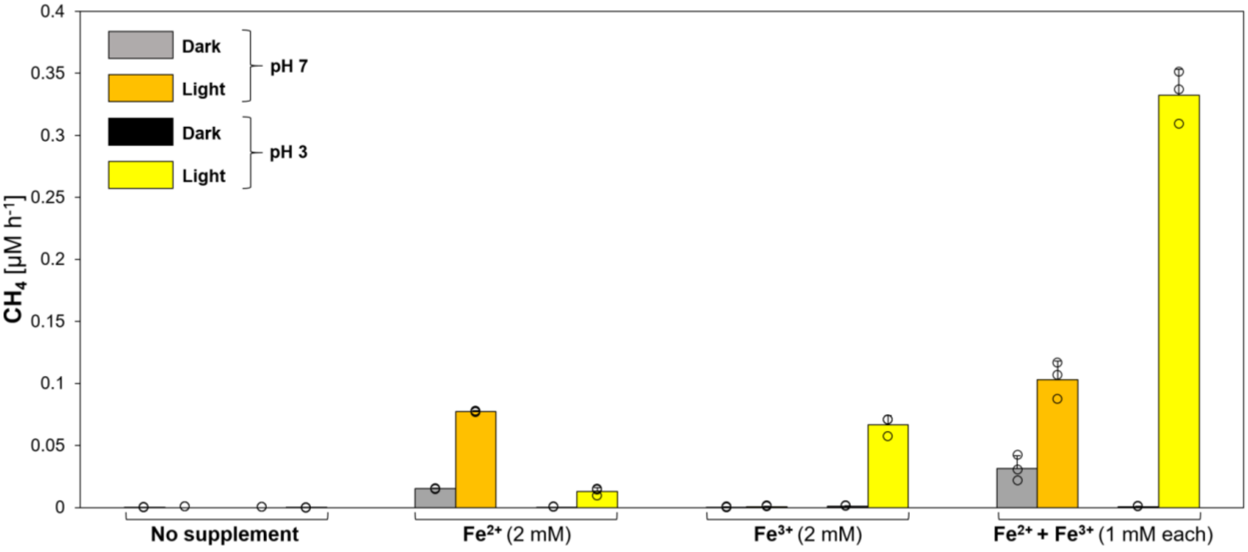


**Fig. S4. Light-driven $CH_4$ formation is enhanced upon $[Fe(H_2O)_6]^{3+}$ photolysis under acidic conditions.** Only trace $CH_4$ levels (<0.0002 µM $h^{-1}$) were measured without iron addition. Upon $Fe^{2+}$-supplementation, $CH_4$ formation rates increased at pH 7 from ~0.015 µM $h^{-1}$ in the dark to ~0.077 µM $h^{-1}$ under light and at pH 3 from <0.0002 µM $h^{-1}$ in the dark to ~0.013 µM $h^{-1}$ under light, suggesting a light-driven ·OH formation from $OH^-$. Conversely, ~0.67 µM $CH_4$ $h^{-1}$ are formed upon $Fe^{3+}$-supplementation in the light under acidic conditions, while only trace amounts of $CH_4$ were detected under pH-neutral conditions or in the dark. At pH 7, a stoichiometric 1:1 $Fe^{3+}$:$Fe^{2+}$ ratio increased $CH_4$ formation rates by ~1.3-fold to ~0.1 µM $h^{-1}$ in comparison to $Fe^{2+}$-supplemented samples, while corresponding $CH_4$ formation rates at pH 3 increased ~25-fold to ~0.316 µM $h^{-1}$. Samples consisting of buffered solutions (pH 3 or 7) were incubated in the dark or under broad spectrum light (Fig. S3) for 24 h under $N_2$. The bars are the mean + standard deviation of three independent measurements, shown as circles.

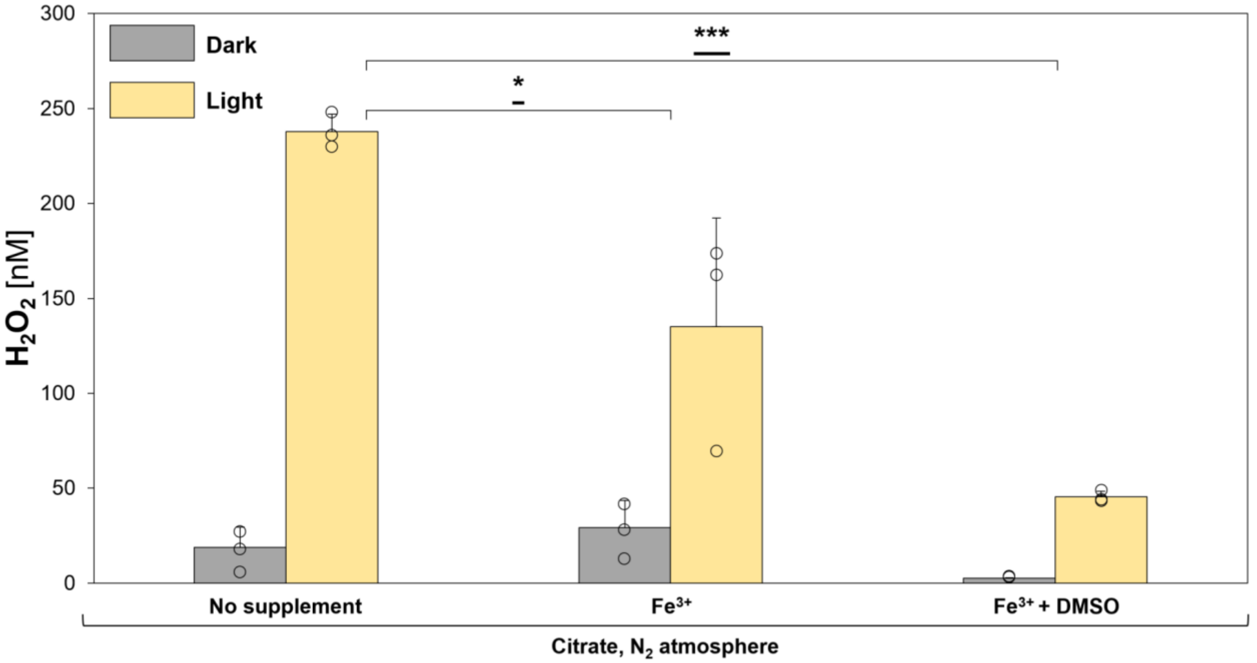


**Fig. S5. Iron and DMSO reduce $H_2O_2$ concentrations generated by light.** Under light, $H_2O_2$ concentrations decrease from ~238 nM (buffer + citrate) to ~135 nM in the presence of iron (buffer + citrate + $Fe^{3+}$). This decrease in $H_2O_2$ concentration indicates the reaction of $H_2O_2$ with $Fe^{2+}$ formed by LMCT. Upon addition of DMSO, $H_2O_2$ levels further decrease to ~45 nM under light, underlining the ROS-scavenging effect of DMSO. In a total volume of 4 mL, samples contained 20 mM potassium phosphate buffer (pH 7), 10 mM citrate and, optionally, 2 mM $FeCl_3$ and 500 mM DMSO that were previously degassed with $N_2$ and kept in an anoxic tent overnight. Samples were incubated in the dark or under illumination (Fig. S3) for 3 h and directly analysed via fluorescence-based $H_2O_2$ endpoint measurements (see Methods). Statistical analysis was performed using paired two-tailed *t*-tests, *: $p \leq 0.05$, ***: $p \leq 0.001$. The bars are the mean + standard deviation of three independent measurements, shown as circles.

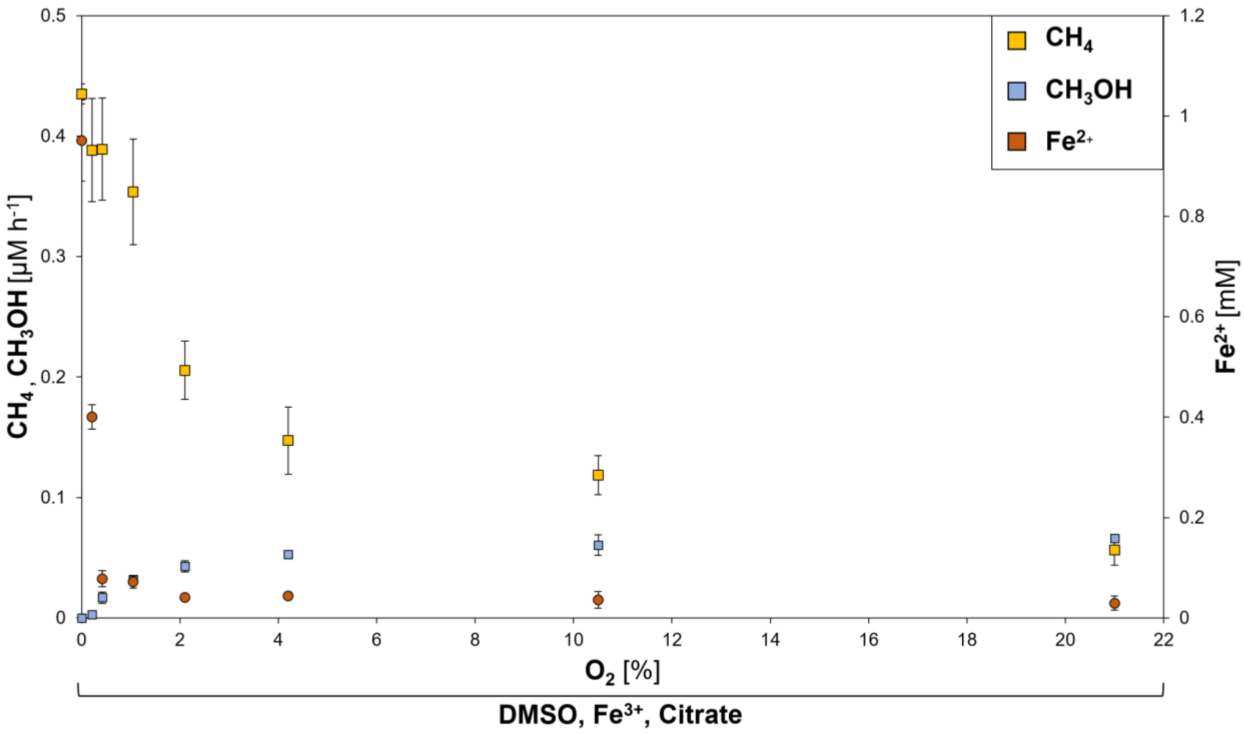


**Fig. S6. Light-driven formation of $CH_4$, $CH_3OH$ and $Fe^{2+}$ from DMSO and $Fe^{3+}$ under varying $O_2$ concentrations.** In a total volume of 4 mL, samples containing 20 mM potassium phosphate buffer (pH 7), 500 mM DMSO, 2 mM $FeCl_3$ and 10 mM citrate were incubated for one day in sealed 20 mL glass vials under constant broad-spectrum illumination (Fig. S3). Mixtures of air and $N_2$ were generated and adjusted to 1 bar, ranging from ~21 % $O_2$ to pure $N_2$ (0 % $O_2$). From 0 % $O_2$ to ~1 % $O_2$, $CH_4$ rates decreased from ~0.44 µM $h^{-1}$ to ~0.35 µM $h^{-1}$, while $Fe^{2+}$ levels decreased from ~1 mM to ~0.07 mM. From ~1 % $O_2$ to 21 % $O_2$, $CH_4$ rates further decreased by an additional ~0.2 µM $h^{-1}$, while $Fe^{2+}$ levels further decreased from ~0.07 mM to ~0.04 mM, indicating an immediate $Fe^{2+}$ oxidation by $O_2$ or the Fenton reaction at high $O_2$ concentrations and a formation of excess $Fe^{2+}$ under suboxic and anoxic conditions. Apart from the anoxic sample (0 % $O_2$), formation of $CH_3OH$ was detected in the presence of $O_2$, ranging from ~0.003 µM $h^{-1}$ (~0.2 % $O_2$) to ~0.07 µM $h^{-1}$ (21 % $O_2$). Data are means ± standard deviation of triplicates.

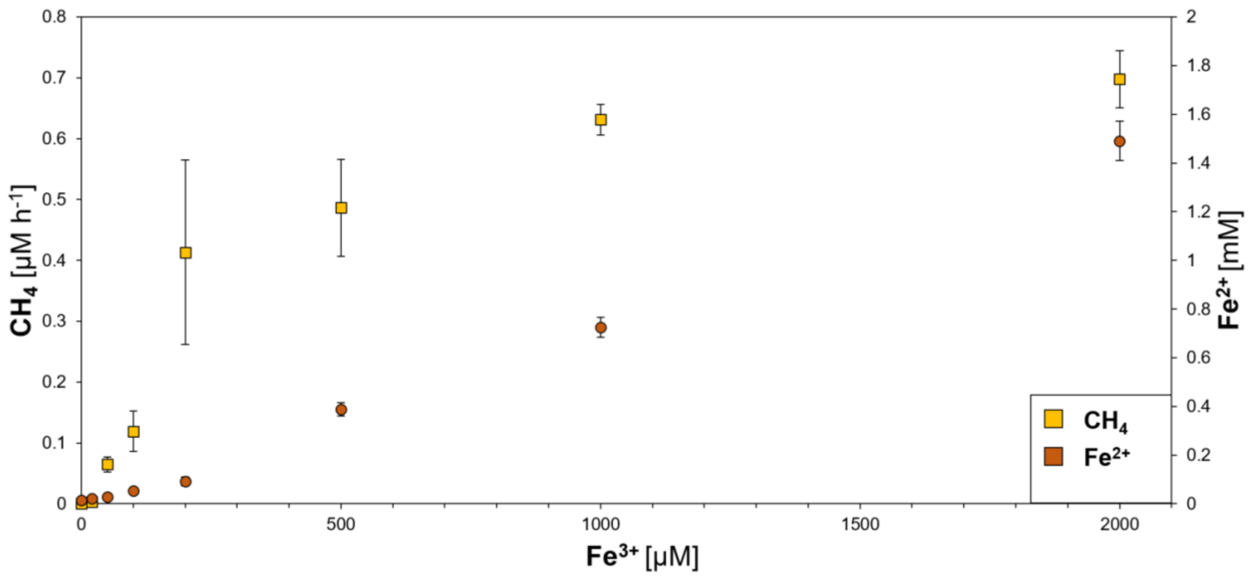


**Fig. S7. Light-driven $CH_4$ formation from DMSO correlates with the iron concentration.** In a total volume of 4 mL, samples containing 20 mM potassium phosphate buffer (pH 7), 500 mM DMSO and 10 mM citrate were incubated for one day under $N_2$ in sealed 20 mL glass vials under constant broad-spectrum illumination (Fig. S3). Different amounts of $Fe^{3+}$ (as $FeCl_3$) were added, ranging from 0 µM to 2000 µM. Only trace amounts of $CH_4$ (<0.01 µM $h^{-1}$) were formed upon supplementation with 0 or 20 µM $Fe^{3+}$. Methane amounts then increased from ~0.06 µM $h^{-1}$ $CH_4$ (50 µM $Fe^{3+}$) to ~0.4 µM $h^{-1}$ (200 µM $Fe^{3+}$), with $Fe^{2+}$ levels also increasing from ~0.03 mM to ~0.11 mM. From 200 µM $Fe^{3+}$ to 2000 µM $Fe^{3+}$ supplementation, $CH_4$ levels increased by ~0.3 µM $h^{-1}$. In the same range, $Fe^{2+}$ levels increased from ~0.11 mM to ~1.5 mM, indicating that LMCT-driven $Fe^{2+}$ formation occurred at higher rates as its consumption to $Fe^{3+}$ via the photochemical Fenton reaction. Data are means ± standard deviation of triplicates.

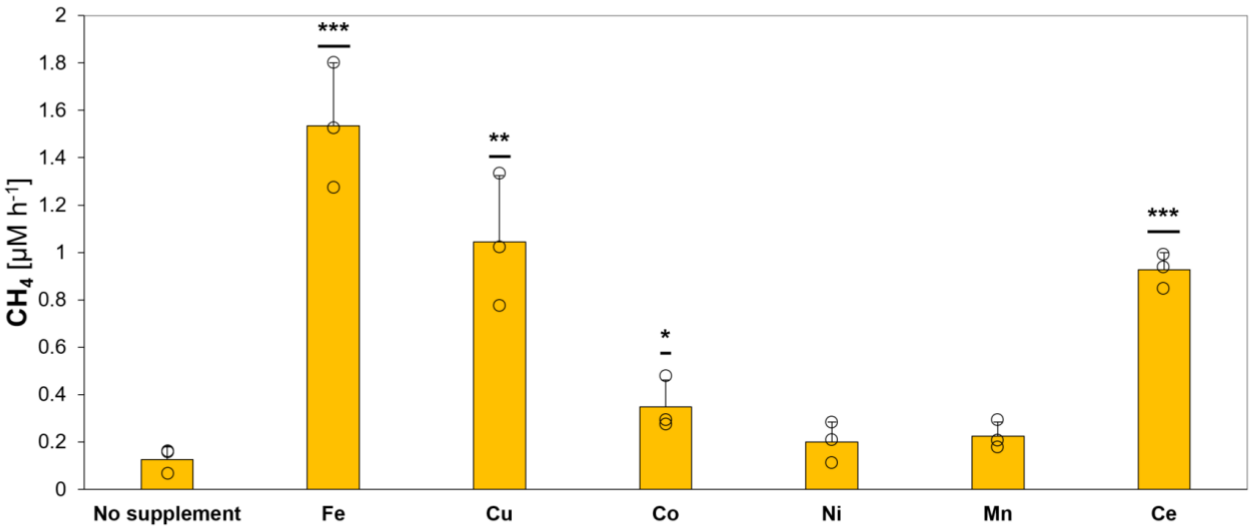


**Fig. S8. Light-driven formation of $CH_4$ from DMSO by reduced transition metals.** In a total volume of 4 mL, samples containing 20 mM potassium phosphate buffer (pH 7), 500 mM DMSO and 10 mM ascorbate were further supplemented with 2 mM cerium ($CeNH_4SO_4$), manganese ($MnSO_4$), cobalt ($CoNO_3$), nickel ($NiSO_4$), copper ($CuCl_2$) or iron ($FeCl_3$) and incubated for one day under $N_2$ in sealed 20 mL glass vials under constant broad-spectra illumination (Fig. S3). Supplementation with iron, copper, cobalt and cerium led to significant increases of observed $CH_4$ amounts, ranging from ~0.35 µM $h^{-1}$ (Co) to ~1.04 µM $h^{-1}$ (Fe). Statistical analysis was performed using paired two-tailed *t*-tests, *: $p \leq 0.05$, **: $p \leq 0.01$, ***: $p \leq 0.001$. The bars are the mean + standard deviation of three independent measurements, shown as circles.

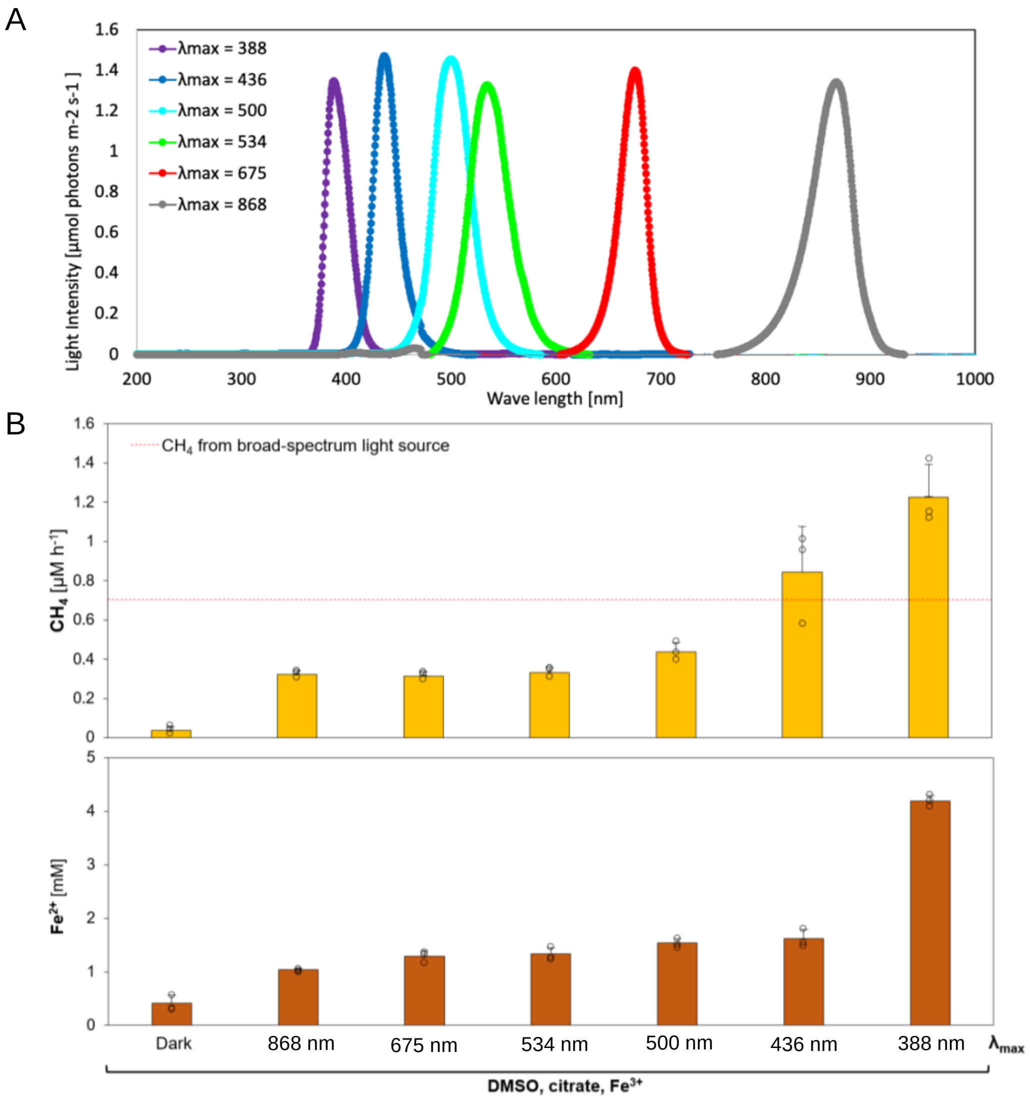


**Fig. S9. Light-driven $CH_4$ formation from DMSO increases in the near-UV spectrum. (A)** LED light spectra. Sample triplicates were illuminated under different light spectra ($\lambda_{max}$ = 388 nm, 52 ± 2 µmol photons $m^{-2}$ $s^{-1}$; 436 nm, 67 ± 2 µmol photons $m^{-2}$ $s^{-1}$; 500 nm, 96 ± 6 µmol photons $m^{-2}$ $s^{-1}$; 534 nm, 95 ± 1 µmol photons $m^{-2}$ $s^{-1}$; 675 nm, 57 ± 6 µmol photons $m^{-2}$ $s^{-1}$; or 868 nm, 103 ± 11 µmol photons $m^{-2}$ $s^{-1}$). Each triplicate was illuminated by one LED light (averaged light spectra presented above). **(B)** In a total volume of 4 mL, samples containing 20 mM potassium phosphate buffer (pH 7), 500 mM DMSO, 5 mM $Fe^{3+}$ (as $FeCl_3$) and 10 mM citrate were incubated under $N_2$ in sealed 20 mL glass vials. For one day, samples were illuminated with LED lights at different wavelengths. ~0.04 µM $h^{-1}$ $CH_4$ and ~0.3 mM $Fe^{2+}$ were formed in the dark. Upon illumination at $\lambda_{max}$ = 868 nm, ~0.3 µM $h^{-1}$ $CH_4$ and ~1 mM $Fe^{2+}$ were measured, increasing to ~0.45 µM $h^{-1}$ $CH_4$ and ~1.6 mM $Fe^{2+}$ at $\lambda_{max}$ = 500 nm. At shorter wavelengths, $CH_4$ amounts further increased to ~0.85 µM $h^{-1}$ ($\lambda_{max}$ = 436 nm) and ~1.2 µM $h^{-1}$ ($\lambda_{max}$ = 388 nm), while $Fe^{2+}$ levels increased up to ~4.3 mM at $\lambda_{max}$ = 388 nm. The dashed red line depicts the average $CH_4$ amounts obtained from samples illuminated by a broad-spectrum light source (Fig. S3, intensity: ~62 µmol $s^{-1}$ $m^{-2}$). The bars are the mean + standard deviation of three independent measurements, shown as circles.

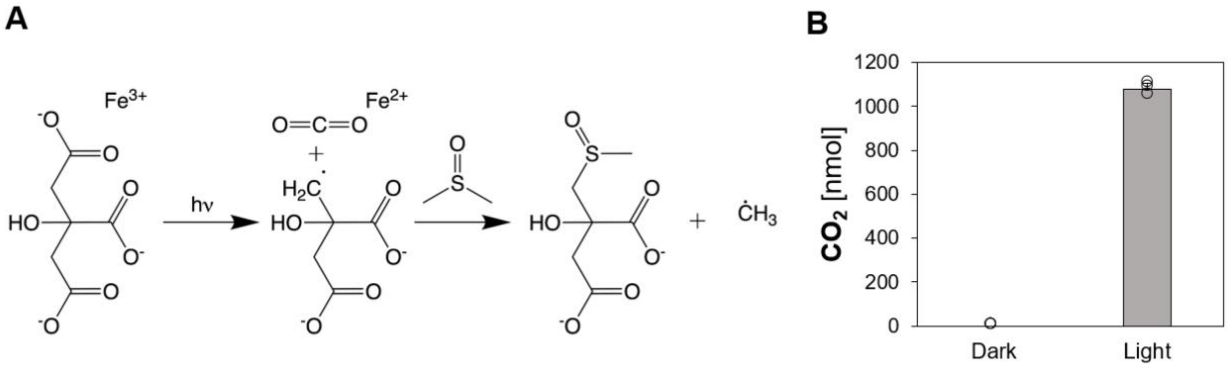


**Fig. S10. Decomposition of the LMCT-induced citrate radical drives ·$CH_3$ formation from DMSO.** **(A)** Illumination of a $Fe^{3+}$-citrate complex results in $Fe^{2+}$ and a citrate radical, disassembling into $CO_2$ and a carbon-centered radical that further reacts with DMSO by cleaving off a ·$CH_3$. **(B)** While no $CO_2$ is formed in the dark, ~1080 nmol $CO_2$ is detected from illuminated samples containing 2 mM $FeCl_3$, 10 mM citrate, 500 mM DMSO in 20 mM potassium phosphate buffer (pH 7) under $N_2$, after 24 h. The bars are the mean + standard deviation of three independent measurements, shown as circles.

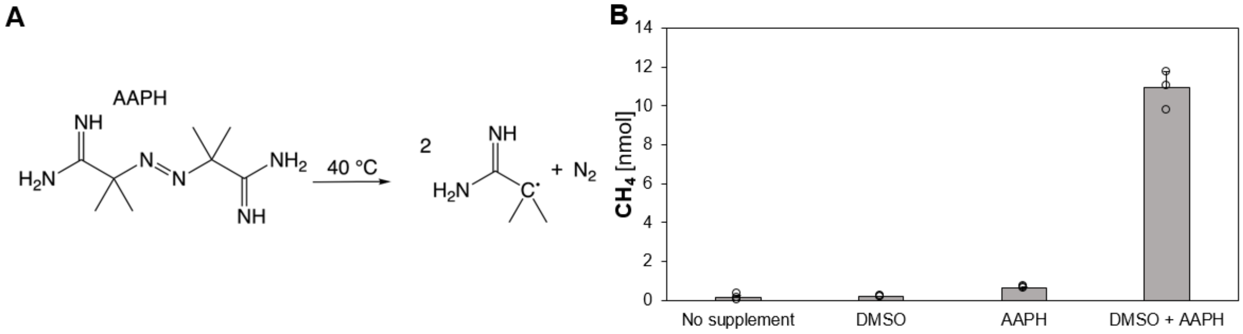


**Fig. S11. Organic radicals drive $CH_4$ formation from DMSO. (A)** At 40 °C, the compound 2,2′-Azobis(2-methylpropionamidine) dihydrochloride (AAPH) decomposes into 2 carbon-centered organic radicals and $N_2$. **(B)** $CH_4$ is formed upon the interaction between DMSO and AAPH. While only ~0.2 nmol or ~0.6 nmol $CH_4$ are formed from DMSO or AAPH alone, respectively, ~11 nmol $CH_4$ are formed upon combining both substances, indicating $CH_4$ formation driven by carbon-centered radicals. 500 mM DMSO was mixed with 20 mM AAPH and incubated for 3 h in $N_2$-saturated DPBS buffer at 40°C under $N_2$.

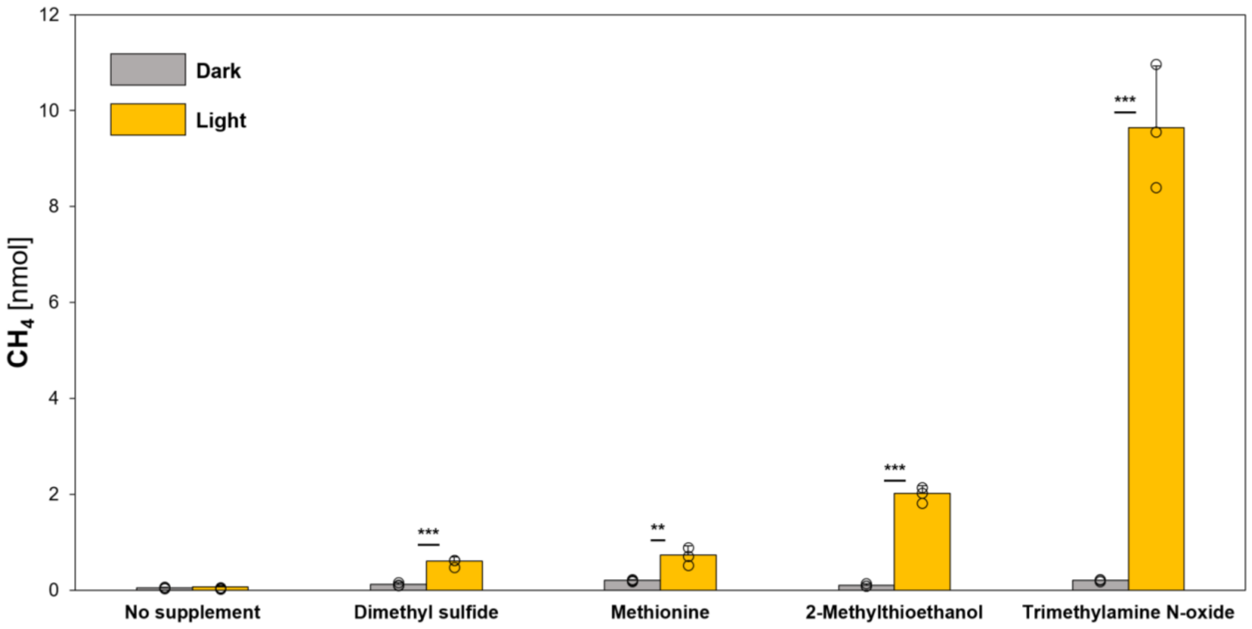


**Fig. S12. Light-driven formation of $CH_4$ from methylated S-/N-compounds.** In a total volume of 10 mL, samples containing 20 mM potassium phosphate buffer (pH 7), 10 mM $FeCl_3$ and 100 mM citrate were supplemented with 500 mM methylated S-/N-compounds and incubated for two days under $N_2$ in sealed 20 mL glass vials at 97 °C. Upon illumination (Fig. S3), significant increases in $CH_4$ levels were measured for dimethyl sulfide, methionine, 2-methylthioethanol and trimethylamine N-oxide, ranging from ~0.61 nmol (dimethyl sulfide) to ~9.65 nmol (trimethylamine N-oxide). Statistical analysis was performed using paired two-tailed *t*-tests, **: $p \leq 0.01$, ***: $p \leq 0.001$. The bars are the mean + standard deviation of three independent measurements, shown as circles.

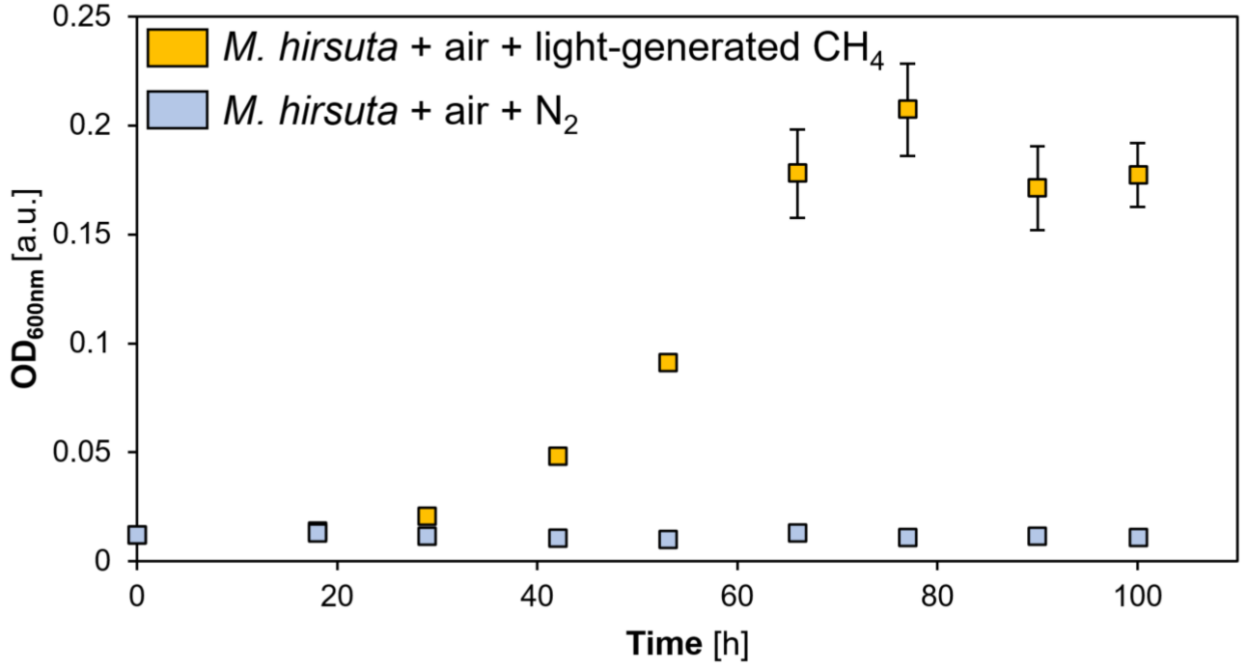


**Fig. S13. Light-driven $CH_4$ formation sustains methanotrophic growth of *Methylocystis hirsuta*.** Methanotrophic growth of *M. hirsuta* is sustained by light-generated $CH_4$. While $CH_4$-supplemented samples (yellow) grew, $N_2$-supplemented samples (blue) did not exhibit bacterial growth (see Methods). Data are means ± standard deviation of three independent measurements.
